# Evaluating the persistence and utility of five wild *Vitis* species in the context of climate change

**DOI:** 10.1101/2021.12.10.472174

**Authors:** Jonas A. Aguirre-Liguori, Abraham Morales-Cruz, Brandon S. Gaut

**Affiliations:** Department of Ecology and Evolutionary Biology University of California, Irvine

**Keywords:** Climate change, Crop Wild Relatives, Genetic offset, Genomic Load, Local adaptation, Migration Load, Species Distribution Models

## Abstract

Crop wild relatives (CWRs) have the capacity to contribute novel traits to agriculture. Given climate change, these contributions may be especially vital for the persistence of perennial crops, because perennials are often clonally propagated and consequently do not evolve rapidly. By studying the landscape genomics of samples from five *Vitis* CWRs (*V. arizonica, V. mustangensis, V. riparia, V. berlandieri* and *V. girdiana*) in the context of projected climate change, we addressed two goals. The first was to assess the relative potential of different CWR accessions to persist in the face of climate change. By integrating species distribution models with adaptive genetic variation, additional genetic features such as genomic load and a phenotype (resistance to Pierce’s Disease), we predicted that accessions from one species (*V. mustangensis*) are particularly well-suited to persist in future climates. The second goal was to identify which CWR accessions may contribute to bioclimatic adaptation for grapevine (*V. vinifera*) cultivation. To do so, we evaluated whether CWR accessions have the allelic capacity to persist if moved to locations where grapevines (*V. vinifera*) are cultivated in the United States. We identified six candidates from *V. mustangensis* and hypothesized that they may prove useful for contributing alleles that may mitigate climate impacts on viticulture. By identifying candidate germplasm, this work takes a conceptual step toward assessing the genomic and bioclimatic characteristics of CWRs.

## INTRODUCTION

Natural populations respond to climate change by adapting to new conditions, by tolerating wider environmental ranges (phenotypic plasticity), by migrating to suitable areas, or by going extinct (Feeley, Rehm, & Machovina, 2012). There is a pressing need to better understand each of these mechanisms to manage and conserve species (Des Roches, Pendleton, Shapiro, & Palkovacs, 2021; Frankham, 2005; Waldvogel et al., 2019). This need is even more pressing for the wild relatives of crop plants (or crop wild relatives, CWRs), because CWRs often contain novel genetic diversity that can contribute to crop improvement (Janzen, Wang, & Hufford, 2019). The potential loss of CWRs has repercussions beyond biodiversity: they are crucial for breeding crops that meet the climate challenge (McCouch, 2013).

Given their importance, numerous studies have focused on the geographic distributions, *ex situ* germplasm collections and *in situ* conservation of CWRs (e.g., (Castañeda-Álvarez et al., 2016)). Surprisingly fewer studies have assessed the potential effects of climate change on CWRs (Jarvis, Lane, & Hijmans, 2008), but recent notable exceptions include studies of CWRs in Europe (Aguirre-Gutiérrez, van Treuren, Hoekstra, & van Hintum, 2017) and the United States (US) (Khoury et al., 2020). These and similar studies typically employ species distribution models (SDMs) to infer the climatic niche of CWRs based on the locations where the species is (or is not) found. This niche is then projected into the future, based on the predicted climate. The outcome of SDMs is an estimate of the predicted geographic niche under climate change, which helps inform the fate of species. SDMs are limited, however, because they usually ignore biotic interactions (Lawler, White, Neilson, & Blaustein, 2006), phenotypic plasticity, and the potential for evolutionary adaptation. As a consequence, SDMs likely yield biased predictions about species persistence (Exposito-Alonso et al., 2018; Hällfors et al., 2016; Ikeda et al., 2017; Razgour et al., 2018)

Although the number of climate-based studies of CWRs is growing, fewer studies have assessed the fate of CWRs by integrating climate predictions with landscape genomic data (Aguirre-Liguori, Ramírez-Barahona, & Gaut, 2021; Aguirre-Liguori, Ramírez-Barahona, Tiffin, & Eguiarte, 2019). Genomic data can be used to model how adaptive genetic variation will change in the context of climate predictions. In this way, one can identify species or populations that appear genetically poised to adapt to climatic change and, conversely, populations that may require especially dramatic genetic changes and are thus genetically endangered. This integration of genomic data with climate predictions is relatively new (Capblancq, Fitzpatrick, Bay, Exposito-Alonso, & Keller, 2020; Fitzpatrick & Keller, 2015), and hence it has been applied to only a handful of taxa (see Capblancq et al., 2020). Studies of poplars, pearl millet and *Arabidopsis* have been especially notable, because they have shown that predicted shifts in allelic variants correlate with components of fitness (Exposito-Alonso, Burbano, Bossdorf, Nielsen, & Weigel, 2019; Fitzpatrick, Chhatre, Soolanayakanahally, & Keller, 2021; Rhoné et al., 2020), thus arguing for the relevance of the approach.

While genomic data include potential information about the adaptive process, the data also contain information about non-adaptive processes that can provide insights into the potential evolutionary fate of species (Aguirre-Liguori et al., 2021; Waldvogel et al., 2019). Such information includes the history of genetic migration among populations, insights into the history and potential effects of genetic drift, and the magnitude of genomic load, which correlates inversely with fitness (Frankham, 2005). To incorporate additional genetic and ecological information into species predictions, Aguirre-Liguori et al. (2021) proposed the FOLDS model, a conceptual framework to evaluate potential responses to climate change. The goal of the FOLDS model is to predict which populations are most likely to respond adequately to climate change, based on multiple layers of information. Those layers can include SDMs, information about putative climate-adaptive alleles, phenotypic data and other population genetic information, such as population size and genomic load.

In this study, we employ the FOLDS model to assess potential responses to climate change for samples from five North American (NA) *Vitis* CWRs. We focus on the CWRs of *Vitis* because they are climatically diverse (Callen, Klein, & Miller, 2016) and agronomically crucial. *Vitis* CWRs are employed for hybrid scion breeding and also as rootstocks, to the extent that ∼80% of viticulture worldwide utilizes rootstocks from NA *Vitis* species (Ollat et al., 2016). Among ∼25 NA *Vitis* species (Wan et al., 2013), those from the American Southwest are of particular interest, because some are resistant to important diseases and others grow under abiotic stresses like extreme conditions that may ‘preadapt’ them to some aspects of climate change (Heinitz, Uretsky, Peterson, Acosta, & Walker, 2019). As concrete examples, *V. berlandieri* is used as a rootstock especially in limestone soils and hot, dry environments (Heinitz et al., 2019), and *V. arizonica* has been used to create hybrid grapevine scions that are resistant to Pierce’s Disease (Summaira Riaz et al., 2009), an economically devastating disease caused by a bacterium that infects several economically important crops (Rapicavoli, Ingel, Blanco-Ulate, Cantu, & Roper, 2018).

We also focus on *Vitis* because climate change is predicted to disrupt the phenology and production of domesticated grapevines (*V. vinifera*) (Morales-castilla et al., 2020), representing substantial impact to a ∼$5 billion per annum industry (USDA, 2013). In addition, despite their ubiquitous use, rootstocks have a remarkably narrow genetic foundation. Currently, a collection of ∼10 rootstocks are used for 90% of grafted grapevines. The most common rootstocks include hybrid or solo contributions from seven NA *Vitis* species. However, the contribution from each species is often (and, in fact, usually) a single accession (Marín et al., 2021). These observations underscore the pressing need to identify additional germplasm for rootstock and scion breeding (Heinitz et al., 2019; Summaira Riaz et al., 2019), given the pressures of climate change.

With genomic data from samples representing five *Vitis* species, this study has two goals. The first is to employ the FOLDs model to evaluate which sets of individuals in the sample are most likely to persist through climate change at their sampled locations in nature. Our evaluation considers climate predictions, the complement of putatively adaptive alleles, resistance to Pierce’s Disease and aspects of population history. The second goal is to assess whether accessions have adaptive alleles that may allow them to persist where *V. vinifera* is currently grown in the United States in predicted future climates. The motivation for this second goal is to evaluate whether CWRs have combinations of alleles that may make them potentially useful, pending extensive future functional validation, for scion or rootstock breeding. Overall, our work takes a conceptual step toward estimating the climate impacts on CWRs and toward identifying candidate species and accessions that may prove useful for the *in situ* adaptation of an important crop.

## METHODS

### Plant material, resequencing and variant identification

The plant material used in this study consisted of 105 individuals from five American *Vitis* species (*V. arizonica, n* = 22; *V. mustangensis, n* = 24; *V. berlandieri, n* = 22; *V. girdiana, n* = 18 and *V. riparia; n* = 19), which were described previously in Morales-Cruz et al. (2021) (Table S1). The 105 accessions were also assayed previously for resistance to *Xylella fastidiosa* (Morales-Cruz et al., 2021; S. Riaz, Huerta-Acosta, Tenscher, & Walker, 2018), the causative agent of PD. Briefly, PD resistance was evaluated using greenhouse experiments in which *X. fastidiosa* was inoculated in different individuals and concentration was evaluated using ELISA tests 10 to 14 weeks after inoculation. Individuals were considered to be resistant to PD if they had concentrations of *X. fastidiosa* < 13 Least Square Means of colony forming units (CFUs) per mL (Summaira Riaz, Tenscher, Heinitz, Huerta-Acosta, & Andrew Walker, 2020).

We used the sequencing data generated by Morales-Cruz *et al*. (2021), which is in the Short Read Archive at NCBI under BioProject ID: PRJNA731597, and their SNP calls. Briefly, they filtered Illumina paired-end reads of 150 base pairs (bp) and mapped them to the *V. arizonica* b40-14 v1.0 genome (Morales-Cruz et al., 2021) (https://doi.org/10.5281/zenodo.4977234 and www.grapegenomics.com/pages/Vari/). Joint SNP calling was conducted using the HaplotypeCaller in the GATK v.4.0 pipeline following (Zhou et al. (2017). The VCF files were split by species, and the raw SNPs were filtered with bcftools v1.9 (https://samtools.github.io/bcftools/) and vcftools v0.1.15 (https://vcftools.github.io/). SNPs sites were kept for downstream analyses if they were biallelic, had no missing data, had quality higher than 30, had a depth of coverage higher than five reads, and also had no more than three times the median coverage depth. Additionally, the following expression was applied under the exclusion argument of the filter function in bcftools: “QD < 2.0 | FS > 60.0 | MQ < 40.0 | MQRankSum < −12.5 | ReadPosRankSum < −8.0 | SOR > 3.0”. These steps resulted in a range of high-quality filtered SNPs from 3.6 million in *V. girdiana* to 5.6 million in *V. riparia* (Table S2).

### Species Distribution Models

For each CWR *Vitis* species and for *V. vinifera* we downloaded occurrences from the Global Biodiversity Information Facility (www.gbif.org; 2020; DOI10.15468/dl.4emr87). We removed duplicated locations and those that were potential misclassifications due to being obvious outliers. We also downloaded 19 bioclimatic variables from Worldclim 2 (Fick & Hijmans, 2017) for a period representing the present, which is an average of observations from 1970 to 2010, and to 54 forecasts of climate change (FCCs). Our rationale for including 54 FCCs was to incorporate uncertainty in climate projections. The forecasts were downloaded at a 2.5 minute resolution from Worldclim 2 (last accessed May 2022), based on the CMIP6 project (Eyring et al., 2016). The 54 FCCs corresponded to five circulation models (GFDL-ESM4, IPSL-CM6A-LR, MPI-ESM1-2-HR, MRI-ESM2-0, UKESM1-0-LL), three time periods (2041-2060 [mean 2050]; 2061-2080 [mean 2070]; 2081-2100 [mean 2090]) and four Shared Socio-economic pathways (SSPs) that model different trajectories of greenhouse effects (SSPs 126, 245, 370, 585). [We were unable to acquire the data for the GFDL-ESM4 model with SSPS 245 and 585 for any time period, resulting in 54 instead of total 60 FCCs.]

To build an SDM for each species, we used previously published scripts (Aguirre-Liguori et al., 2021) that performed four steps. First, the script removed correlated bioclimatic variables (r>0.8) with the highest Variance Inflation Factor, to moderate multicollinearity between variables. Second, it identified a calibration or background area, which corresponded to the migration layer in the BAM (Biotic, Abiotic and Migration) model (Peterson et al., 2011; Soberón, 2010) and to terrestrial eco-regions of the world (Olson et al., 2001). Third, the script employed the BIOMOD2 package in R (Thuiller, Georges, Engler, Georges, & Thuiller, 2016) and the Maxent algorithm (SJ Phillips, Anderson, & Schapire, 2006; Steven Phillips & Dudík, 2008) to build and project the models. Finally, we performed 20 bootstrap replicates per model. For each replicate we used 70% of occurrences to train the model and 30% to test the model with True Skill Statistics for tenfold internal cross-validation.

Based on the present-day SDMs, we projected the potential distribution of species to the 54 FCCs. For each species and each FCC we estimated how the geographic distribution (i.e., the number of pixels in the projection) was expected to change from the present. Ultimately, we evaluated the mean change and the dispersion across the FCCs in each of the three projected time-periods. We also estimated whether each of the 105 accessions was expected to persist in a given FCC. That is, for each individual at a given time period, we counted how many times FCCs predicted the persistence of the individual’s sample location. If the location was predicted to be outside the species’ estimated niche for all 20 FCC models at a specific time-period, we coded that population as extinct by that time period and into the future.

### Investigating patterns of local adaptation

#### Identification of outlier SNPs

For each CWR species sample, we used Baypass (Gautier, 2015) to identify outlier loci that had significant associations with the first four principal components (PC) of bioclimatic variables (Figure S1). [Figure S1 provides a schematic overview of methods that incorporated genomic and environmental data, as applied to the FOLDS model.] The PCs were determined with the *prcomp* function in R, based on the 19 bioclimatic variables from the present-day. Baypass analyzes patterns of covariation among accessions to account for population structure and then analyzes the correlation between a variable (in this case each PC) and the allelic frequencies of individual SNPs across individuals. A SNP was deemed an outlier if it had a significant correlation with a variable that was not explained by the covariance between populations. Following Jeffreys’ rule (Gautier, 2015), we characterized outlier SNPs as those that had a Bayes Factor (BF) > 20.

#### Local Genomic Offsets

To calculate genetic offsets, we used gradient forest (GF; Figure S1) (Ellis, Smith, & Roland Pitcher, 2012), a machine learning method that models the turnover in genetic composition across the landscape (Fitzpatrick & Keller, 2015). GF identifies which bioclimatic variables contribute importantly to the construction of the model, the SNPs that are associated with bioclimatic variables, and the adaptive genetic composition across the predicted climate (Capblancq et al., 2020; Fitzpatrick & Keller, 2015; Waldvogel et al., 2019). When applied to bioclimatic data from both the present and the future, GF also estimates the local genetic offset (or ‘local offset’), an estimate of the amount of genetic change necessary to adapt in the future at the same locality (Capblancq et al., 2020; Fitzpatrick & Keller, 2015). The local offset depends on the genetic composition of populations and the magnitude of the environmental change that is expected to occur; populations with higher offsets are expected to be more vulnerable to climate change (Fitzpatrick & Keller, 2015).

For each species, we obtained the GF model using the *gradient forest* package in R (Ellis et al., 2012) for all the outlier SNPs identified by Baypass (**Table 1**), based on allelic frequencies within individuals (Figure S1). These frequencies were either 0 or 1 for homozygotes or 0.5 for inferred heterozygotes. The local offset was then calculated as the Euclidian distance of the genetic compositions between the present and different time periods (Fitzpatrick & Keller, 2015) (Figure S1). We used the GF models to predict how genetic compositions were expected to change in each of the 54 FCCs.

**TABLE 1.** Five *Vitis* CWR samples, with summary statistics.

| Species | N<sub>ind</sub><sup>1</sup> | Ne (95% CI)<sup>2</sup> | N<sub>Bay</sub><sup>3</sup> | N<sub>GF</sub><sup>4</sup> | BIO<sup>5</sup> |
| --- | --- | --- | --- | --- | --- |
| *V. arizonica* | 22 | 35,448 (33,658-37,237) | 13,531 | 5,792 | 3,15,2,9 |
| *V. berlandieri* | 22 | 40,112 (37,008-43,215) | 8,425 | 3,371 | 4,7,3,10 |
| *V. mustangensis* | 24 | 74,839 (59,722-89,957) | 8,787 | 2,730 | 18,5,10,4 |
| *V. girdiana* | 18 | 53,666 (47,065-60,267) | 5,650 | 5,254 | 4,7,3,11 |
| *V. riparia* | 19 | 49,279 (45,454-53,105) | 10,626 | 8,720 | 10,4,8,1 |

<sup>1</sup> N<sub>ind</sub> is the number of samples per species
<sup>2</sup> the mean effective population size estimated across 6 samples with 95% confidence intervals
<sup>3</sup> N<sub>Bay</sub> refers to the number of outlier SNPs identified with Baypass
<sup>4</sup> N<sub>GF</sub> refers to the number of outlier SNPs identified with Gradient Forest
<sup>5</sup> BIO includes the list of the four top bioclimatic variables identified with GF, in decreasing order of importance

GF is traditionally applied to allele frequencies across populations, but here we applied it to individual genotypes. To investigate whether results were applicable for individuals, we performed two tests. First, we transformed allelic frequencies within individuals to pseudo-population frequencies, using a probabilistic model, and reanalyzed the data (see Supplemental Methods). Our simulations suggest that the sampling of individuals, instead of populations, had little impact on the construction of the GF model, the estimation of genomic offsets and the identification of important bioclimatic variables (See **Supplementary Methods** and **Figure S2**). Second, we used a population dataset that had been used to calculate genetic offsets previously (Aguirre-Liguori et al. 2021) and evaluated the effect of subsampling from 1 to 11 individuals per population on the estimation of local offsets (see Supplementary Methods). We found that using individuals instead of allelic frequencies reduced the estimated local offsets across populations but also that the rank correlation among offsets was >0.99. This high correlation indicated that the relative magnitude and order of local offsets remained similar across locations and sampling strategies (Figure S3), further suggesting that use of individuals in GF is a suitable for our comparisons.

#### Calculating the Adaptive Score

In addition to local offsets, we calculated other parameters from the data to help summarize genetic information that can be used to evaluate climate persistence. For example, to summarize information about adaptive alleles we calculated an adaptive score (*S_a_*). *S_a_* measured the proportion of alleles across all candidate SNPs that were ‘pre-adapted’ to climate, based on climate projections (Figure S1). As used here, *S_a_* is modelled after the population adaptive index (PAI) of Bonin et al. (2007). The PAI measured the proportion of loci that have a selective signal within populations – i.e., the proportion of loci that were outliers due to high allelic frequency deviations among populations. In our case, we identified the alleles that -- according to turnover functions from GF analyses -- were predicted to be adaptive in future climates (Figure S4).

To identify pre-adapted alleles, we assessed the projected direction of environmental change for important bioclimatic variables (i.e., change across the *x*-axis in **Figure S4**) by identifying ‘pre-adapted’ individuals that, according to turnover functions, require small change in their genetic composition in the future (i.e., small changes across the y-axis, **Figure S4**). After identifying pre-adapted individual(s), for each candidate SNP we built a linear model (using *lm* in R) to confirm that the genotypic state correlated significantly with the expected bioclimatic variable. Significantly correlated alleles were deemed adaptive, and the high frequency allele was categorized as the adaptive state. Given adaptive alleles, we calculated *S_a_* as the total number adaptive alleles per individual across all candidate SNP sites, divided by 2x the number of candidate SNPs (The factor of 2x reflected the diploid state). Thus *S_a_* reflected the proportion of adaptive alleles that were present in an individual across all candidate SNPs. It ranged from 0 to 1, where 0 indicates an individual with no adaptive alleles and 1 indicates homozygosity of adaptive alleles at all candidate SNPs.

For each species, we calculated *S_a_* for each of the four bioclimatic variables that had the strongest contribution to building the GF model and then calculated the mean across the four variables (Figure S1). Further details, tests for potential biases and rationale for *S_a_* are provided in the **Supplementary Methods**. We note that *S_a_* overlaps with, but differs from, local offsets in two ways. First, high local offsets can be caused either by strong predicted environmental shifts or by extensive expected genetic shifts. In contrast, *S_a_* focuses solely on genetic composition by counting whether an inferred adaptive allele is present at each position. Second, local offsets cannot be compared across species (Láruson, Fitzpatrick, Keller, Haller, & Lotterhos, 2022), but *S_a_* can because it reports a proportion – i.e., of adaptive alleles across candidate climate-related SNPs.

### Additional Population Genetic Parameters

#### Demographic history

To incorporate information about genetic drift into the FOLDS model, we estimated the effective population (*Ne*) size of each species using MSMC2 v2.1.1 (Mallick et al., 2016) with unphased SNPs. To include only the most informative genomic regions, we created a mappability mask and a coverage mask. We created the mappability mask for the *V. arizonica* b40-14 v1.0 genome using the software SNPable (http://lh3lh3.users.sourceforge.net/snpable.shtml). We generated 150 bp mers moving in 1 bp increments across the genome, mapped the sequences back to the genome with BWA v0.7.8-r455 (Li & Durbin, 2009) and identified “mappable” genomic regions where the majority of sequences mapped uniquely without mismatches. To include only the regions with sufficient sequencing coverage, we calculated the coverage from the alignment file of each sample with the bedcov program from the samtools (v1.10) package, supplying a bed file of 10 kb non-overlapping windows across the genome. We then created a mask per sample that only included regions with sequencing coverage higher than 5x. We then used unphased SNPs from the six samples per species with the highest average coverage as input for the run in MSMC2. For analyses, we assumed a mutation rate of 5.4e-9 (Liang et al., 2019) and a generation time of 3 years (Zhou et al., 2017). In addition to MSMC2 with unphased data, we applied MSMC2 to phased data and also used SMC++ as a another source for *Ne* estimation (see Supplementary Methods). The results of the most recent time index from SMC++ and MSC2-unphased were highly similar (**Figure S5**). The results were also very consistent among individual MSMC2 runs (**Figure S6),** and so we showed the average per species (**Figure S5**).

#### Genomic load

Genomic load is a measure of the number of predicted deleterious alleles in an individual. To identify deleterious alleles, we first determined the functional context (exonic, intronic, and intergenic) of SNPs based on *V. arizonica* B40-14 genome annotations ((Morales-Cruz et al., 2021), https://doi.org/10.5281/zenodo.4977234 and www.grapegenomics.com/pages/Vari/). Exonic SNPs were annotated to be synonymous, nonsynonymous or frameshift mutations using SnpEff version 5.0e (Cingolani et al., 2012). Nonsynonymous SNPs were predicted as deleterious or tolerated using the SIFT score (Ng & Henikoff, 2003), as computed in the program SIFT 4G (Vaser, Adusumalli, Leng, Sikic, & Ng, 2016). SIFT scores ≤ 0.05 were interpreted as deleterious and SIFT scores > 0.05 were considered to be tolerant. Because the reference genome biases the distribution of missing data (Lohmueller et al., 2008), we only used polymorphic sites without missing data that were predicted across all five wild grape species, resulting in a set of 397,723 SNPs.

### Applying the FOLDS model

To assess the potential effects of climate change, we employed the FOLDS model (Aguirre-Liguori et al., 2021), based on six layers of genomic, phenotypic and climate information described above (Figure S1). We used UpsetR (Conway, Lex, & Gehlenborg, 2017) to plot intersections among the six layers of information for each of the 54 FCCs. For each UpsetR model, we evaluated which accessions passed specific filters for the six layers. As described below (see Results), we selected different thresholds and filters for inclusion and synthesis. For example, accessions were included based on SDMs if the sampling location was predicted to be in the species’ niche in the future and also when they were resistant to *X. fastidiosa* (i.e., had concentrations < 13 Least Square Means of CFUs/mL). We also selected accessions as persistent if they had local offsets and genomic load values below 90%, 50% and 25% of their distribution across species. Since local offsets are not comparable between species (Láruson et al., 2022), the thresholds for local offsets were considered only within species. Finally, accessions were included when they had *S_a_* or *Ne* above the 10%, 50% and 75% of the distributions across species.

### Migration Load

The FOLDS model defined a set of CWR accessions that were best situated to persist in the face of climate change. We focused on these accessions to evaluate their genetic offset at locations where *V. vinifera* is currently cultivated in the United States. This measure of offset, which has been called either the “forward offset” (Gougherty, Keller, & Fitzpatrick, 2021) or the “migration load” (Rhoné et al., 2021), estimates whether an accession has the allelic complement to persist when moved to a different location. Thus, the migration load is the genetic offset for an individual calculated between its present-day sampling site and for predicted climates at other locations. In this case, the other locations represent current locations of viticulture. The migration load is a tool to evaluate whether CWR accessions might be useful in future climates where grapevines are currently cultivated.

We estimated migration load for accessions that passed the FOLDs threshold for each of the 54 FCC models at the 1278 United States locations of *V. vinifera* cultivation available from gbif.org. Following previous work (Aguirre-Liguori et al., 2021; Rhoné et al., 2020), the migration load was calculated as the Euclidian distance of the genetic composition between the present location of an accession and the future estimate in the projected climate of *V. vinifera* cultivation. We identified locations where the migration load was within the current range of the species’ predicted local offsets over the same FCC period. Finally, for each *V. vinifera* location, we counted how many times an accession had a migration load within the range of its local offsets. We interpreted these accessions as candidates for having climate-related alleles that could be useful to viticulture in the face of climate change.

## RESULTS

### Species Sampling and Data

In this study, we used previously published resequencing data from 105 individuals representing five North American *Vitis* species: *V. arizonica*, *V. mustangensis* (synonym *V. candicans*), *V. berlandieri*, *V. girdiana* and *V. riparia* (Morales-Cruz et al., 2021). These species are either currently used as rootstocks, represent the parents of hybrid rootstocks and scions, or have phenotypic properties that make them potentially valuable to viticulture (Heinitz et al., 2019). The two most closely related species in this study (*V. girdiana* and *V. arizonica*) split at least 10 million years ago (Wan et al., 2013) and perhaps even earlier (Morales-Cruz et al., 2021), so they have not diverged especially recently.

For each of the five species, 18 to 24 accessions were resequenced (**Table 1**). We employed the previously determined SNPs (Morales-Cruz et al., 2021), which were used previously to explore the divergence dynamics among species and their history of hybridization. Here, however, we focused on a subset of SNPs that had no missing data within each species. The resulting number of SNPs ranged from 3.6 million in *V. girdiana* to 5.6 million in *V. riparia* (**Table S2**). The accessions, their sampling locations, and their SNPs provided the basis for evaluating species’ persistence under projected climate change.

### Evaluating the Persistence of Wild *Vitis* Under Predicted Climate Change

To assess the potential effects of climate change, we combined six layers of genomic and climate information within the FOLDS framework (Aguirre-Liguori et al., 2021). The six layers were chosen to include different types of information, including: i) SDMs to incorporate niche information; ii) patterns of local adaptation and predicted allelic responses to climate change; iii) insights into genetic drift and genomic load, both of which may affect evolvability and persistence; and iv) phenotypic characterizations of resistance to PD. Below we describe each of the six layers before synthesizing information within FOLDS framework.

#### Layer 1 - Species distribution models

We estimated SDMs to examine expected changes in the geographic distribution of the five wild *Vitis* species over 54 climate models (FCCs), focusing on four time periods (present-day, 2050, 2070 and 2090), five circulation models and four shared socio-economic pathways (see Methods). For each of the five species, we projected the SDM onto the 54 FCCs. Then, for each SDM we measured the projected habitable area of each species (**Figure 1a**), as has been done previously for wild Vitis (Callen et al. 2016; Heibnitz et al, 2019) but also the predicted trends over time (**Figure 1b**). Strikingly, the average amount of habitable area was predicted to increase over time for all species, but with a marginal reduction for *V. girdiana* after 2050 (**Figure 1b, Table S3**).

**FIGURE 1.**
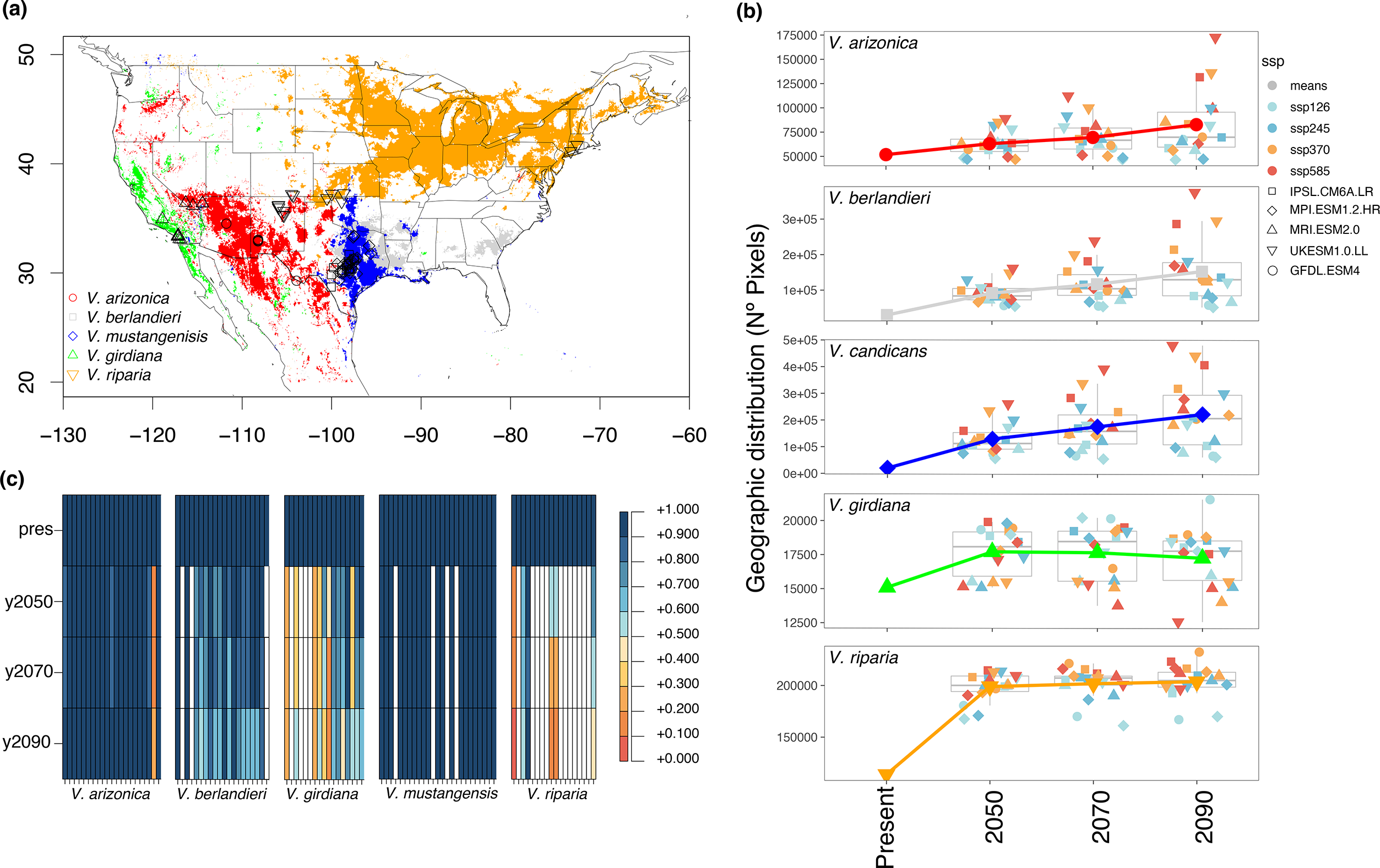
Results based on SDMs of the five *Vitis* species in this study. (a) Present-day SDMs showing the estimated bioclimatic niche of each of the five species. Species are colored according to the key. The map also provides the sampling locations of the 105 accessions, with shapes and colors of different species provided by the key. (b) The geographic area of the niche for each species graphed into the future for 54 future climate change (FCC) models and their mean. The symbols and colors represent the estimated geographic size for different circulation models and shared socio-economic pathways, respectively. The central line and symbols across the years 2050, 2070 and 2090, represents the mean of SDMs based on the different FCC models. The colors represent the species, as indicated in the key of panel A. (c) Each column represents one of the 105 accessions in this study. The color in the column indicates the proportion of times that an accession was predicted to exist for a given time period across the 18 FCC models. If the location of sampled individual did not fall within the future niche for all FCC models for a given time period, as defined by the SDM, then it is no longer represented in the figure (white column). Hence, by 2090, SDMs predict that most *V. arizonica* and *V. mustangensis* samples will be able to persist in their current locations, as will a few *V. berlandieri*. In contrast, most *V. riparia* in our sample are not predicted to persist.

Although the overall expansion of niche area is promising, there was substantial variation in the details across the 54 FCCs. For example, the dispersion across models increased as a function of time, and the SSPs that predicted higher greenhouse gas emissions predicted larger CWR distributions (**Figure 1b**). Overall, the SDMs predicted many *V. arizonica*, *V. mustangenis* and *V. berlandieri* locations were likely to persist until 2090 (**Figure 1c, Table S3**). However, persistence varied substantially among species, because from 20% to 43% of sampling localities (depending on the species and climate models), were predicted to disappear from species’ niches by 2090. Overall, SDMs predict that distribution of the wild *Vitis* species is expected to increase in the future, but the results also suggest that many populations represented by the samples in our study will need to either adapt or migrate to survive.

#### Layer 2 - Local Offsets and Adaptative Variation

We used GF to estimate the local offsets of *Vitis* individuals within each species. Following common practice (Capblancq et al., 2020; Fitzpatrick, Keller, & Lotterhos, 2018), we ran GF with a subset of outlier SNPs that were associated with environmental variables. To identify these outlier SNPs, we performed genome environment associations (GEA) with Baypass (Gautier, 2015), identifying SNPs correlated to the first four principal components (PCs) of 19 present-day bioclimatic variables (Figure S1). [The first 4 principal components explained 99.9% of bioclimatic data for *arizonica;* 97.3% for *berlandieri* 97.3%; 92.9% for *mustangensis*; 98.8% for *girdiana* 98.8% and 97.4% for *riparia 97.4%*]. Our GEA detected from 5,650 to 13,531 outlier SNPs within each species (**Table 1**). We applied GF to the set of outlier SNPs from the GEA, yielding a subset of 2,730 to 8,720 candidate SNPs within each species that had significant non-linear associations across samples (**Table 1**), based on present-day bioclimatic data.

GF analyses provided three additional outcomes (Figure S1). The first was a set of bioclimatic variables that were important for building the model, as measured by the contribution of the variable to the construction of the model (R^2^> 0). For simplicity, we focused on the four most important variables for each species, some of which (**Table 1**) were shared across species. For example, BIO10 (the mean temperature of the warmest quarter) had a strong contribution to the building of GF models in *V. berlandieri, V. mustangensis* and *V. riparia,* whereas BIO4 (temperature seasonality) was a top contributor for all species but *V. arizonica* (**Table 1**).

The second outcome was estimates of local genetic offsets, which we computed for each individual in each species based on contrasts between present-day climate and each of 54 FCC models (see Methods), resulting in 5,670 (= 54 models x 105 individuals) estimates. We compared rank correlations among offsets between all FCC models to determine if there was a consistent offset signal. Pairwise comparison revealed that >95% of pairs had rank correlations of *rho*>0.5, suggesting that different climatic models generally showed similar offset trends (Figure S7). For simplicity, in Figure 2a we show the genetic offsets in each species for a single model (IPSL-CM6A-LR_ssp585_2061-2080), because this model had the highest average rank correlation (mean rho = 0.83) compared to the remaining 53 models. The corresponding results for the remaining models are presented in Table S1.

**FIGURE 2.**
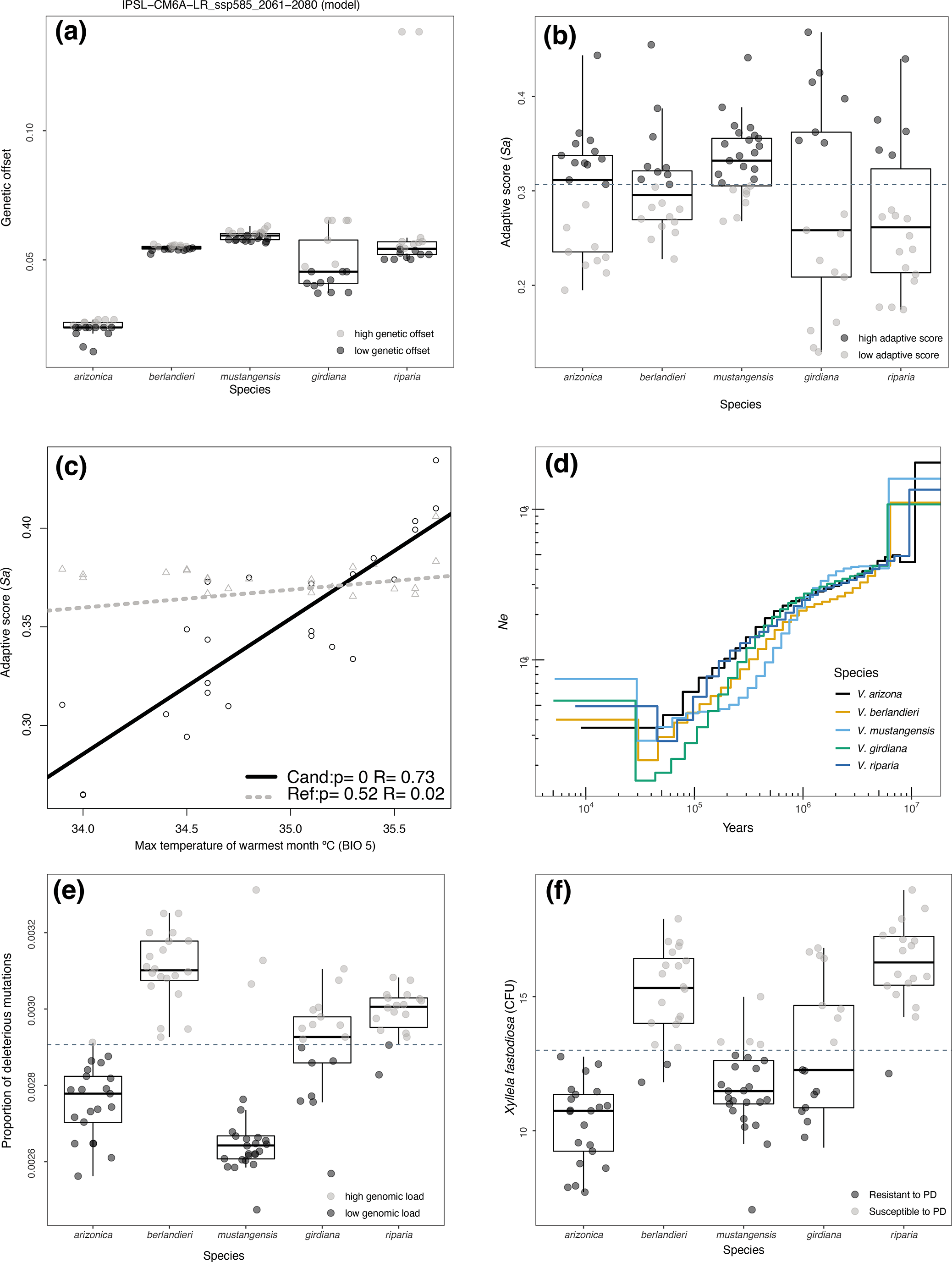
Layers of information used in the FOLDS model. (a) Distribution of local genomic offset for each species for future climate change (FCC) model IPSL−CM6A−LR_ssp585_2061−2080. We darkened circles based on their status above and below the median within each species. (b) Variation in the adaptive scores (*S_a_*) across species. Each circle represents a sampled accession. The dashed line shows the median of *S_a_* across all accessions; circles are darkened if they have *S_a_* above the median. (c) An example of correlations between *S_a_* and bioclimatic variables, in this case the maximum temperature of the warmest month (BIO 5) for *V. mustangensis* samples. The dashed line shows the correlation for reference SNPs. For all results and test for potential biases see **Figure S8.** (d) Change in effective population size across time for each species. The color of each species is indicated by the key. (e) The measure of genomic load – i.e., the proportion of predicted deleterious SNPs per genome across a shared set of SNPs – plotted for each accession and species. [Note that the proportion of deleterious SNPs is equivalent to the number of deleterious SNPs per genome, since all inferences were based on a shared set of SNPs.] The dashed line represents the median of the genomic load. Darker circles are below the median. (f) Distribution of the concentration of *X. fastidiosa* in different wild *Vitis* species after exposure to the bacterium. The dashed line represents the value 13 CFUs, which has been the boundary between resistance and susceptibility. Darker circles represent accessions with CFUs < 13.

Local offsets cannot be compared quantitatively across species (Láruson et al. 2022), because they are estimated with different SNPs and bioclimatic information. They do, however, help predict whether specific alleles are expected to be adaptive in future climates. To build a metric for cross-species comparison, we utilized local offsets to inform a measure, *S_a_*, that measures the proportion of adaptive alleles within each individual and that can be compared across individuals and species (see Methods). For each species, we calculated the mean adaptive score (*S_a_*) (see Methods & Figure S1). Like offsets, *S_a_* varied within species (**Figure 2b**, **Table S1**); comparing *S_a_* across species indicated that our *V. girdiana* and *V. riparia* samples have pre-adaptive alleles in lower proportions than the other species (**Figure 2b**).

Because *S_a_* had not been applied previously, we performed tests to evaluate whether *S_a_* was biased by geographic, demographic or climatic variables. As expected, *S_a_* was strongly and significantly correlated with at least one of the four most important bioclimatic variables identified by GF (**Figure 2c**); in fact, it was correlated with all four for three of 5 species (**Figure S8**). As expected, *S_a_* was more highly correlated to bioclimatic variables than a similar score (*S_ref_*) calculated from a set of reference SNPs (**Figure S8;** see Methods), suggesting it contains additional information. The notable exception was for *V. girdiana* samples, where *S_ref_* was strongly correlated with all four of the top 4 associated bioclimatic variables (**Figure S8**), likely reflecting genetic structure (Morales-Cruz et al., 2021) or other features that complicate interpretation of *S_a_* in this species.

#### Layer 4- Effective population size

The efficacy of selection depends on adaptive diversity but also on population factors, like genetic drift and genomic load, that are typically ignored in climate studies (Aguirre-Liguori et al., 2021; Brady et al., 2019; Nadeau & Urban, 2019). To assess the potential influence of genetic drift, we estimated historical effective population sizes (*N_e_*) over time, using MSMC2 (Schiffels & Wang, 2020). For all species we found that *N_e_* decreased from the distant past until ∼20,000 years ago, followed by recent moderate recoveries (**Figure 2d**). From these analyses, we retrieved the current estimate of *N_e_*, which ranged from ∼35,000 individuals in *arizonica* to ∼75,000 in *V. mustangensis* (**Table 1**); *V. mustangensis* also yielded the highest *N_e_* estimate based on another method, SMC++ (Terhorst, Kamm, & Song, 2017) (**Figure S5**) and the results were consistent among individual MSMC2 runs (**Figure S6**). The relatively high *N_e_* for *V. mustangensis* was surprising, because it currently has a narrow geographic distribution (**Figure 1a**), but it nonetheless suggests that it may be less prone to drift effects than other species.

#### Layer 5 - Genomic Load

Genomic load, the accumulation of deleterious mutations in a population, can predict fitness (Frankham, 2005). We estimated genomic load within and between species by measuring the number of potentially deleterious alleles per accession (see Methods). To account for differences in the number of SNPs between species and potential reference biases, we focused only on SNPs shared between species, with no missing data. Overall, genomic load indicated that *V. mustangensis*, *V. arizonica* and *V. riparia* individuals had relatively low genomic load among species, but there was also notable variation among individuals within species (**Figure 2e**).

#### Layer 6 - Resistance to Pierce’s Disease

Biotic interactions between species can play a key role in shaping population’s responses to climate (Bascompte, García, Ortega, Rezende, & Pironon, 2019; Zamora-Gutiérrez, Rivera-Villanueva, Martínez Balvanera, Castro-Castro, & Aguirre-Gutiérrez, 2021). As a case study for including such information, we included data about resistance to the bacterium *X. fastidiosa* (Morales-Cruz et al., 2021; Summaira Riaz et al., 2020). We plotted CFU counts for each individual, revealing variation in PD resistance among individuals and species (**Figure 2f**) – e.g., 100% of *V. arizonica* and 87% of *V. mustangensis* accessions were PD resistant while the *V. girdiana* distribution was notably bimodal, with ∼50% of accessions resistant (CFU < 13). In contrast, only one *V. riparia* and two *V. berlandieri* accessions were resistant.

### The FOLDS framework

Ideally, assessing the response of populations to climate change should be based on multiple layers of information (Razgour et al., 2018; Waldvogel et al., 2019); we relied on the preceding six layers to surmise whether an accession is relatively well situated to adapt to climate change. Such accessions should have low offsets, low genomic loads, high *S_a_* values, high *N_e_* (low drift), PD resistance and predicted persistence based on SDMs.

We applied the FOLDS (Aguirre-Liguori et al., 2021) using different thresholds (Figure S1). For example, we first filtered accessions based on all six layers, with thresholds defined by: PD resistance (CFU < 13), SDM persistence, among the lowest 25% of local offset and genomic load within a species, and among the highest 25% of Ne and Sa within a species. We illustrated the intersection of layers for the IPSL-CM6A-LR_ssp585_2061-2080 climatic model (Figure 3a), based on the set of summary statistics (Table S4). The figure shows some of the combinations of layers, and the number of accessions that pass that combination of filters. We can see, for example, that five accessions passed all six layers, suggesting that they are best poised to contribute alleles to adapt climate change, based on location, climate and genomic data. In addition, results for the model in Figure 3a, we provided the complete set of summary statistics used for the UpsetR analyses, with results for the additional 53 climate models (Table S1). These additional tables can be used to determine which accessions are suited to respond adequately to climate change for a particular FCC model.

**FIGURE 3.**
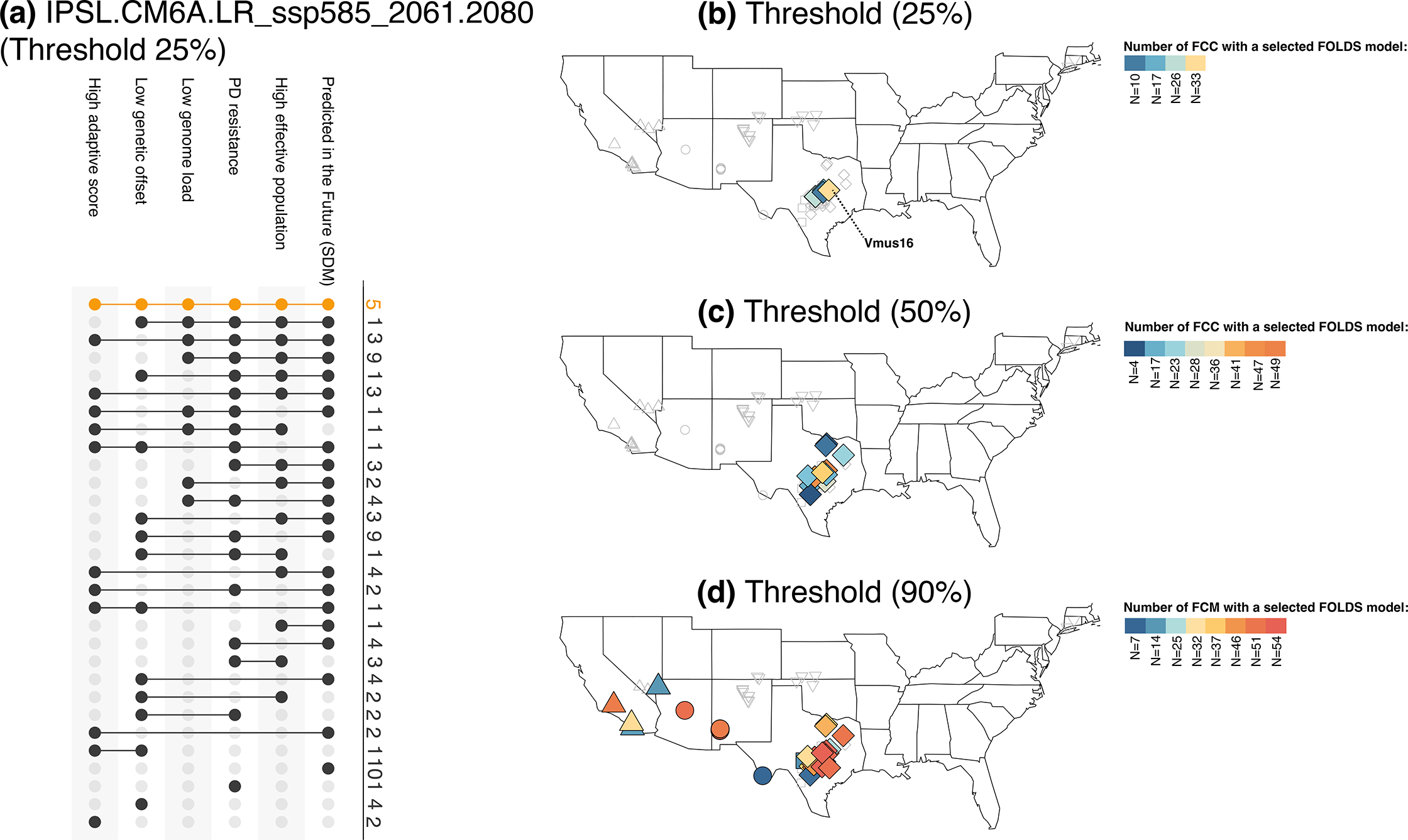
The FOLDS model applied to wild *Vitis*. (a) The upsetR plot shows the intersection across all 105 individuals that pass thresholds (at the strictest 25% threshold) for the six layers of information and for future climate change model IPSL−CM6A−LR_ssp585_2061−2080. Black dots joined by a line indicate that individuals pass the threshold for those layers of information. The number of individuals presenting different intersections are at the right side of the plot. The orange line at the top, indicated that 5 individuals pass the thresholds for 6 layers of information. (b-d) The maps plot the individuals that remained after applying each of the three thresholds (25%, 50%, 90%). The color of the symbols indicate the number of times an individual passes the 6 layers of information based on the 54 FCC models. Table S7 reports the number of times an individual passed the FOLDS test for different thresholds FCC models.

Since we examined 54 FCCs, we calculated the number of times an accession passed the threshold for all six layers for each FCC model (Figure 3b-c). Overall, we found that for the conservative thresholds (0.25 and 0.5), only accessions of *V. mustangensis* passed all six filters for at least one climate model (**Figure 3b-c**). For the 25% threshold, four *V. mustangensis* accessions (Vmus02, Vmus03, Vmus14 and Vmus16) passed all filters for at least 48% of the 54 climate models. Based on this information, we predict that *V. mustangensis* accessions are relatively well situated to persist to predicted shifts in climate or to contribute adaptive variation in the future.

We also relaxed the thresholds, so that (for example) accessions remained when they retained among the lowest 90% of offset and genomic load within a species, and among the highest 10% of *N_e_* and *S_a_* within a species (**Figure 3c-d**). At these threshold levels, a few accessions of *V. girdiana* and *V. arizonica* passed the filters. Importantly, the exercise also provided potential insight into relative risks among taxa. For example, only one *V. riparia* accession remained, even after applying 90% thresholds, for most models (Figure 3d). These results suggest that this species is particularly susceptible to population extinction within our sampled areas.

### Assessing the potential of wild *Vitis* to mitigate climate change in viticulture

Given insights into the potential persistence of wild species, our next task was to evaluate their potential to aid viticulture in the face of climate change. To evaluate which CWRs might be best suited for *in situ* adaptation of the crop, we relied on the concept of migration load (Rhoné et al. 2021), which is the genetic offset for an individual calculated between its present-day sampling site and other locations. In this case, the ‘other locations’ were all the 1278 *V. vinifera* locations in the United States identified in GBIF (**Figure 4a, Table S5**).

**FIGURE 4.**
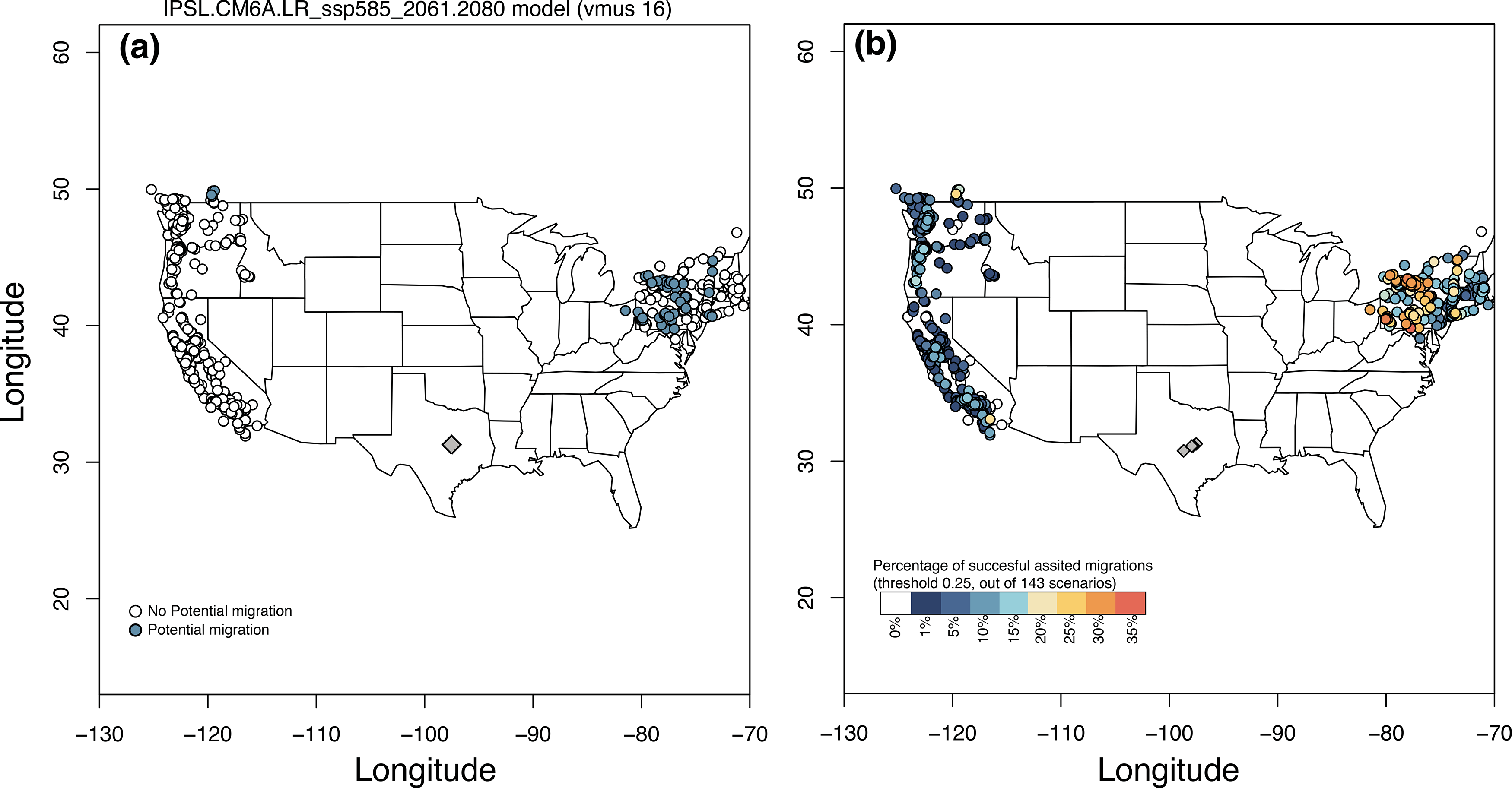
Climate change and in situ adaptation of *V. vinifera*. (a) An example of the locations where the migration load of *Vmus16* (see Figure 3) would be higher (white circles) or lower (blue circles) than the local genetic offset of *V. mustangesis* if it was moved to locations where *V. vinifera* currently grow. This was performed using the future climate change (FCC) model IPSL−CM6A−LR_ssp585_2061−2080. *Vmus16* would be more likely to be a suitable candidate for *in situ* adaptation of grapevine in the NE of the distribution. (b) percentage of times that the migration load for 6 *V. mustangensis* individuals identified by the FOLDS model at a 0.25 threshold (Figure 3b) would be lower than the local genetic offset of *V. mustangensis* across the 54 (FCC) models. The colors of the symbols indicate, from all the possible events, the percentage of times than assisted migration could potentially be used for *in situ* adaptation (out of 143 possible events).

The approach and results are perhaps most easily represented by examples. *V. mustangenesis* accession vmus16 was one of six accessions that passed the FOLDS criteria at the strictest threshold level (Figure 3). As an illustration, we calculated the migration load for this accession in one FCC model (IPSL-CM6A-LR_ssp585_2061-2080), (Figure 4a). We then asked: which, if any, of the migration loads fell within the range of local offsets for the wild species? If the migration offset fell within the range of local offsets, we surmised that vmus16 is a potential bioclimatic fit for crop adaptation at that location. Altogether, vmus16 had low migration offsets in many locations of the Northeastern USA and in one location in the Northwest for this single FCC model (**Figure 4a**). These are, then, candidate regions for the use of this CWR, where it may have climate alleles that could contribute to agricultural applications.

We extended these analyses to the six *V. mustangenesis* accessions that passed the filters of the FOLDs model (Figure 4b), calculating the migration load for each accession and each FCC model at all 1278 *V. vinifera* locations. After assessing whether the migration offset fell within the range of local offsets for each FCC model, we counted the proportion of times the *V. mustangenesis* accessions passed the assessment at each location. The resulting map illustrated the potential suitability of *V. mustangensis* at each location (**Figure 4b**). Overall, *V. mustangenensis* was projected to be more suitable for predicted climates in the Northeast US than in the Western US.

We expanded these analyses to all 48 accessions that passed filters for all six layers of the FOLDS model across the three threshold levels. We found 2630 instances in which an accession passed the FOLDS model for at least one FCC model and one threshold level, including 2096 cases for 90% FOLDS threshold, 391 cases for the 50% threshold, and 143 cases for strictest (25%) threshold (**Figure 3b-d**). From these, we estimated 2,630 x 1,278=3,361,140 migration loads. Tables S5 & S6 provide the calculated migrations loads and indicates whether they fell within the range of local offsets. These Tables thus provide a preliminary evaluation of the potential climate adaptability of wild *Vitis* accessions in regions of *V. vinifera* cultivation.

## DISCUSSION

Climate change is a threat to all biodiversity (Parmesan, 2006; Parmesan & Yohe, 2003), but the fate of CWRs is particularly important. A growing literature has predicted the fate of CWRs under climate change, based on SDMs, but without explicit consideration of evolutionary change, evolutionary history, or phenotypes. Here we have focused on five *Vitis* CWRs as exemplars to address three goals. The first has been to incorporate genomic data to assess the relative potential of CWRs to persist under climate change. The second has been to assess which CWRs, if any, have bioclimatic and genomic profiles that are of potential agronomic interest, pending further experimental evaluation. Finally, we offer this work as a framework to apply to other CWRs and crop systems.

### Evaluating CWR Persistence: insights and caveats

Predicting how populations and species will respond to climate change is not a simple task because multiple factors --- including evolutionary, human, and ecological processes –impact the response of populations to climate change (Waldvogel et al., 2019). We have combined individual measures, including SDMs and measures of genomic variability, to assess the relative ability of CWRs to respond to climate change. To our knowledge, no formal statistical framework exists to perform this task; instead, we relied on the existing FOLDS framework (Aguirre-Liguori et al., 2021).

We recognize that this approach is not complete for many reasons. For example, myriad additional biotic and abiotic phenomena likely affect the ability of populations to evolve in the face of climate change, but not all have been (or can be) measured. In theory, an advantage of the FOLDS approach is that it can incorporate unlimited layers of information, including additional biotic interactions, genetic and environmental data, and potential human impact data (e.g., urbanization or human-mediated fire threats) (Aguirre-Liguori et al. 2021). However, a corresponding shortcoming is that individual layers are weighted equally and treated independently, which may be inaccurate. For example, we have found that *N_e_* and genomic load are correlated across NA *Vitis* accessions (*r* = −l0.41, p<0.001; **Figure S9**). While this correlation is not unexpected -- because selection is less efficient in small populations -- it implies that information from the two layers is partially redundant. We have also unexpectedly found that genomic load and PD resistance are correlated (*r*=-0.26; *p*< 0.01; Figure S9). Perhaps the simplest explanation for the correlation is that less-fit plants (as measured by genomic load) are more susceptible to disease, but it again demonstrates that two seemingly unrelated layers may not provide fully independent information. One important extension of the FOLDS (or similar) models will be the ability to weight the contribution of individual layers to maximize predictive power. This may not be possible, however, without experimental data from climate experiments that can empirically test the accuracies of predictions. Such experiments are an obvious and important next step.

Additional shortcomings of our analyses relate to our treatment of the *Vitis* dataset. For example, local offsets are typically calculated on estimated allele frequencies within populations (Fitzpatrick & Keller, 2015), but we have applied it to diploid frequencies within individuals. While this may decrease predictive power, and is likely conservative for that reason, both simulations and resampling analyses suggest that our conclusions based on single genotypes are robust (See **Supplementary Methods**). We have also ignored the potential for natural migration, which may also impact opportunities to adapt to climate change.

Nonetheless, our synthesis provides a framework to begin to evaluate the relative risk among CWRs to predicted climate change that goes beyond SDMs and niche modeling. By considering evolutionary information, we predict that *V. mustangensis* accessions are the most apt to persist in the face of predicted climate change. The *V. mustangensis* accessions have good persistence in SDMs, high *Ne*, low genomic load, low genomic offsets and PD resistance. The framework also provides additional information for conserving other wild *Vitis* species. For example, some individuals scored well for five of six layers but have either low *Ne* or low adaptive scores **(Figure 3a**). Based on this information, it may be necessary to aid management of source populations by adding genetic diversity from different populations or species. Finally, we accentuate again that any predictions should, and can, be validated experimentally (Kardos & Shafer, 2018) – e.g., by growing suitable wild *Vitis* accessions in common gardens that simulate climate change and testing for fitness effects.

### Crop adaptation and climate change

Given genomic information from CWRs, we have used the GF framework to assess the potential of specific accessions to grow in the projected climates where domesticated grapevines are currently grown in the United States. The motivation for this approach is that viticulture is under climate risk. For example, Morales-Castilla et al. (2020) have used phenological data to estimate that 56% of current, worldwide growing locations will be lost under 2C° of warming, although this value decreased to 24% when they considered intraspecific variety in phenology among grapevine cultivars. Similarly, Hannah et al. (2013) projected a 25% to 73% decrease in suitable land for viticulture by 2050 among major wine producing regions. Hence, we view our analyses as a step toward assessing whether CWRs have alleles that can help mediate the effects of climate change on viticulture.

An important question is how best to counteract climate threats to crops. Generally, there are three non-exclusive strategies. One is human-assisted migration that shift the locations of cultivation for a specific crop (Sloat et al., 2020). A second is to develop new agricultural regions – i.e., to expand arable land. However, land availability is limited, and the ecological and economic costs of this strategy is especially large (Fita, Rodríguez-Burruezo, Boscaiu, Prohens, & Vicente, 2015). The third strategy is *in situ* adaptation – i.e., breeding cultivars that tolerate stresses associated with climate change in their current locations of cultivation (Sloat et al., 2020). This is a strategy that has been advocated previously for *Vitis* rootstocks (Callen et al, 2016) and one for which genomic data may be helpful. In fact, Genomic data have been used to inform this strategy, as illustrated in a landmark study of pearl millet (Rhoné et al., 2020). In the study, Rhoné et al. (2020) generated genomic data from landraces across the African continent, identified geographic regions where cultivation was at risk due to climate change, and then determined the migration load to determine which landraces have genomic profiles that may thrive if moved to at-risk regions.

Using a similar approach, we have asked which of our sampled CWRs have genetic and bioclimatic profiles that may make them suitable for growing in viticulture locations within the United States. Our CWR sample from the Southwestern U.S. may be particularly suitable for this study, because the accessions are adapted to warm and dry environments and hence may be needed for future breeding programs (Hietnitz et al. 2019). Since the FOLDS model predicted that six *V. mustangensis* accessions are particularly likely to persist under climate change, we focused on estimating their migration loads across a range of climate models. Migration load was often within the range of local offsets for that species (**Figure 4**). Based on these results, we hypothesize that some of these wild accessions may prove useful for *in situ* adaptation of cultivated grapevine and specifically that *V. mustangensis* may prove vital agronomically.

It is important, however, to discuss this hypothesis critically. First, we recognize that *V. mustangensis* is not currently used as a rootstock or for scion breeding. In fact, it may not even be the best candidate CWR for *in situ* adaptation at vulnerable sites, given that we have studied only five of 30 species and 105 accessions. In this context, it is worth noting that *V. riparia* is commonly used as a rootstock but none of the *V. riparia* accessions in our sample fared particularly well in the FOLDS model. We suspect this reflects the fact that our sample is taken from the geographic limits of the species, where genomic load is often higher and fitness may be lower (Willi et al. 2018).

Second, our analyses are clearly only a first step, because many additional features of wild germplasm must be characterized before assessing agronomic utility. For example, although our study is rare for including one potential biotic interactor, there are many more other potential biotic interactors (Griggs, Steenwerth, Mills, Cantu, & Bokulich, 2021) -- especially resistance to phylloxera -- that are crucial for agricultural use. We also are not considering important complexities about viticulture, like irrigation and soil type. It is also an open question whether *in situ* adaption to climate change in grapevine will be better achieved by identifying new rootstocks or by using CWRs for scion breeding. The latter has already been shown to be effective for discrete traits that are governed by a few major loci, such as disease resistance. Indeed, PD resistance at the *Pdr1* locus (Krivanek, Riaz, & Walker, 2006) has been backcrossed into various varieties to introduce PD resistance. Bioclimatic adaptation is polygenic, however, as evidenced by the thousands of SNPs associated with bioclimatic variables in each of these species (**Table 1**) and by the fact that associated SNPs are distributed throughout the chromosomes (Morales-Cruz et al., 2021). Given, the polygenic nature of bioclimatic adaptation, we suspect that *Vitis* CWRs will typically be more effective as rootstocks. That said, rootstocks can have a multitude of effects on the scion, including yield, phenology and drought tolerance that may also display environmental interactions (Warschefsky et al., 2016). Moreover, potential rootstock and scion combinations can be incompatible, for reasons that are not yet well understood genetically (Gaut, Miller, & Seymour, 2019).

Finally, we have assumed that the climate models are accurate. Although we see no way around this assumption, we have tried to represent uncertainty in climate predictions by including 54 FCC models representing different time scales, circulation models and SSPs. Some accessions have low migration loads in some locations across most climate models, providing some consistency to the hope that these can be sources for climate-related alleles that may prove useful. Altogether, we have applied a unique combination of both climate and genomic information to identify potential and unexpected candidates for priority evaluation. We believe that this methodological approximation could be valuable for identifying CWRs of other crop for conservation or improvement in the future.

## ACKNOWLEDGEMENTS

The authors thanks Yongfeng Zhou, Summaira Riaz, Andrew Walker and Dario Cantu for sharing data from previous manuscripts. This work was improved by the comments of three anonymous reviewers and was supported by National Science Foundation grant no. 1741627 to B.S.G. J. A-L was supported in part by a UC-Mexus postdoctoral fellowship.

## DATA ACCESSIBILITY STATEMENT

Sequence data that supports the findings of this study were downloaded from the Short Read Archive at NCBI under BioProject ID: PRJNA731597.

## AUTHOR CONTRIBUTIONS

All authors contributed to the ideas of the study; J.A-L. & A.M-C. performed analyses; all authors wrote the manuscript.

## SUPPORTING INFORMATION

### Supplemental Methods

#### Adding random variation to allele frequencies to evaluate the application of GF to individuals

GF has been applied using allelic frequencies from populations instead of individual genotypes. We evaluated how sampling individuals may impact GF results. To do so we simulated “pseudo-frequencies” by changing the frequency (Δ) of each individual genotype (0+Δ, 0.5±Δ or 1-Δ) in each accession. The deviation Δ in each SNP and each individual was obtained by randomly sampling a value that ranged from 0 to 1, with a beta distribution and means 0.1 and 0.3 (**Figure S2a**). The idea of using two means was to increase the magnitude of Δ to account for a stronger bias in the sampling of individuals. For each pseudo-frequency dataset, we ran the GF model and estimated the 6 strongest BIO variable predicted by the model (R^2^>0), plotted the turnover function for the BIO with the strongest R^2^ and estimated the genetic offset of the pseudo-populations in 2070. These simulations were performed for all species 100 times.

Figure S2 shows the results for all simulations in *mustangensis,* but we found quantitatively similar results for all species. First, we found that generating datasets with pseudo-frequencies did not affect the general patterns of the turnover functions (**Figure S2b**). However, as expected, we did find variance between replicates and also that the power was lower for the simulations with Δ distributions with mean 0.3 than 0.1. Second, we found that in general the four most important variables identified for the real dataset were also found with the simulated datasets (**Figure S2c**). However, as Δ increased we found lower consistency. Importantly, however, the BIO variables that were predicted for the simulated data corresponded to the top 6 variables identified in the real dataset. Finally, when we compared the distributions of the simulated genetic offsets with the real genetic offsets, we found similar patterns (**Figure S2d**), indicating that populations with low offsets in the real dataset also had low offsets in the simulated datasets. While we cannot know the real frequencies in the populations, our results suggest that sampling individuals instead of populations may have a small effect on the estimation of the GF models, and that the general patterns are retained.

#### Testing the effect subsampling different number of individuals to estimate the genetic offsets

We evaluated if subsampling different number of individuals could modify the genetic offset rank between populations. For this, we downloaded the dataset of *Zea mays* ssp. *mexicana* that was used by Aguirre-Liguori et al. (2021) to propose the FOLDS model. We used this dataset because it consists of 23 populations, with between 11 and 15 individuals per population. To test the effect of sampling fewer individuals we wrote a script that 1) subsamples 1 to 11 random individuals from each of the population, 2) performs an outlier analysis (LFMM2); 3) performs GF, 4) estimates the genetic offsets of subsampled populations, 5) calculates the rank correlation between the genetic offsets of the subsampled populations and the genetic offsets of the populations using all the sampled individuals. This was done for each subsampling N1 to N11, 1,000 times. Figure S3 shows the boxplots showing the range of the *r* values of the rank correlations for N1 to N11. We found that for all subsamples the rank correlation between the genetic offsets of the subsampled populations and entire populations was higher than 0.995, strongly suggesting that sampling one individual per population does not dramatically affect the rank order of the population’s genetic offsets. In our study we define the persistence of populations based on thresholds that rely on relative rankings; based on these results, sampling individuals to estimate the genetic offsets may not heavily bias our results.

#### Calculating and Evaluating Sa

We calculated the adaptive score (*S_a_*), which was motivated by an analogous measure, the population adaptive index (Bonin, Nicole, Pompanon, Miaud, & Taberlet, 2007). However, instead of analyzing the proportion of adaptive SNPs in a populations, like the adaptive index, *S_a_* estimates the proportion of alleles that will be adaptive in the future. To find adaptive alleles, we first identified pre-adapted individuals and then ran a linear model for each candidate SNP across all individuals, to evaluate if the genotypic state correlated significantly with bioclimatic variables. If the linear model was not significant, we discarded the SNP. If it was significant, we considered the allelic state that was frequent in pre-adapted individuals to be the adaptive allele. For each individual, we summed the number of pre-adaptive alleles present in the accession (2 for each adaptive allele in the homozygous state, 1 for the heterozygous state) across all candidate loci, and then divided that number by the maximum number of potential adaptive alleles. The maximum number of potential adaptive alleles was calculated as 2 x N_Sp_, where N_Sp_ is the number of candidate loci for a given species. We called this value the adaptive score (*S_a_*), which ranges from 0 to 1, where 0 indicates that the population does not have adaptive alleles and 1 indicates that all their alleles are adaptive. For each species we estimated the linear association between the adaptive score across individuals and the 4 bioclimatic variables that have a strong impact on the GF models. Since the correlation between *S_a_* were in general high (r>0.7), we estimated the mean *S_a_* across individuals for the 4 bioclimatic variables.

#### Demographic inference

In the main text, we reported MSMC2 results based on unphased SNPs, but we also calculated demography based on phased SNPs. To phase, we extracted phase-informative SNPs (PIRs) for each sample as described in Delaneau et al. (2013) (Delaneau, Howie, Cox, Zagury, & Marchini, 2013), which identifies reads that span at least 2 heterozygous SNPs, and used the PIRs and the VCF files for each species by chromosome as input to shapeit2 v2.r904 (Delaneau, Zagury, & Marchini, 2013) to phase the SNPs. We used the “generate_multihetsep.py” script from the MSMC tools (https://github.com/stschiff/msmc-tools) to create the input files for MSMC2 taking into account the mappability mask and the coverage masks. For the phased-SNPs runs, we extracted the phased SNPs of the two individuals with the highest coverage per species to run MSMC2 (**Figure S5**).

We also used SMC++ v1.15.4.dev18+gca077da (Terhorst, Kamm, & Song, 2017). To create a low coverage mask, we calculated sequencing coverage from the alignment files using the bedcov program from the samtools (v.1.10) package, supplying a bed file of 10 Kb non-overlapping windows across the genome. We calculated the sequencing coverage per species by summing the coverage of all individuals in the same genomic window. We classified regions as low coverage per species when they had a coverage lower than 5x. The low coverage masks were supplied together with the VCF files to the *vcf2smc* program of SMC++ for species and each chromosome. The demographic history was calculated per species using the *estimate* program from SMC++.

Comparing among methods, MSMC2 with phased and unphased SNPs yielded similar demographic trajectories over time, with the exception of current *Ne*s, which were up to two orders of magnitude higher with phased data. Since these estimates were much higher than the other two results (i.e., MSMC2 with unphased data and SMC++), we believe errors in the phasing process led to overestimation of Ne. In contrast to the two MSMC2-based analyses, the SMC++ results suggested that each taxon reached their lowest Ne at different time periods (**Figure S5**).

### Supplemental Figures

**Figure S1.**
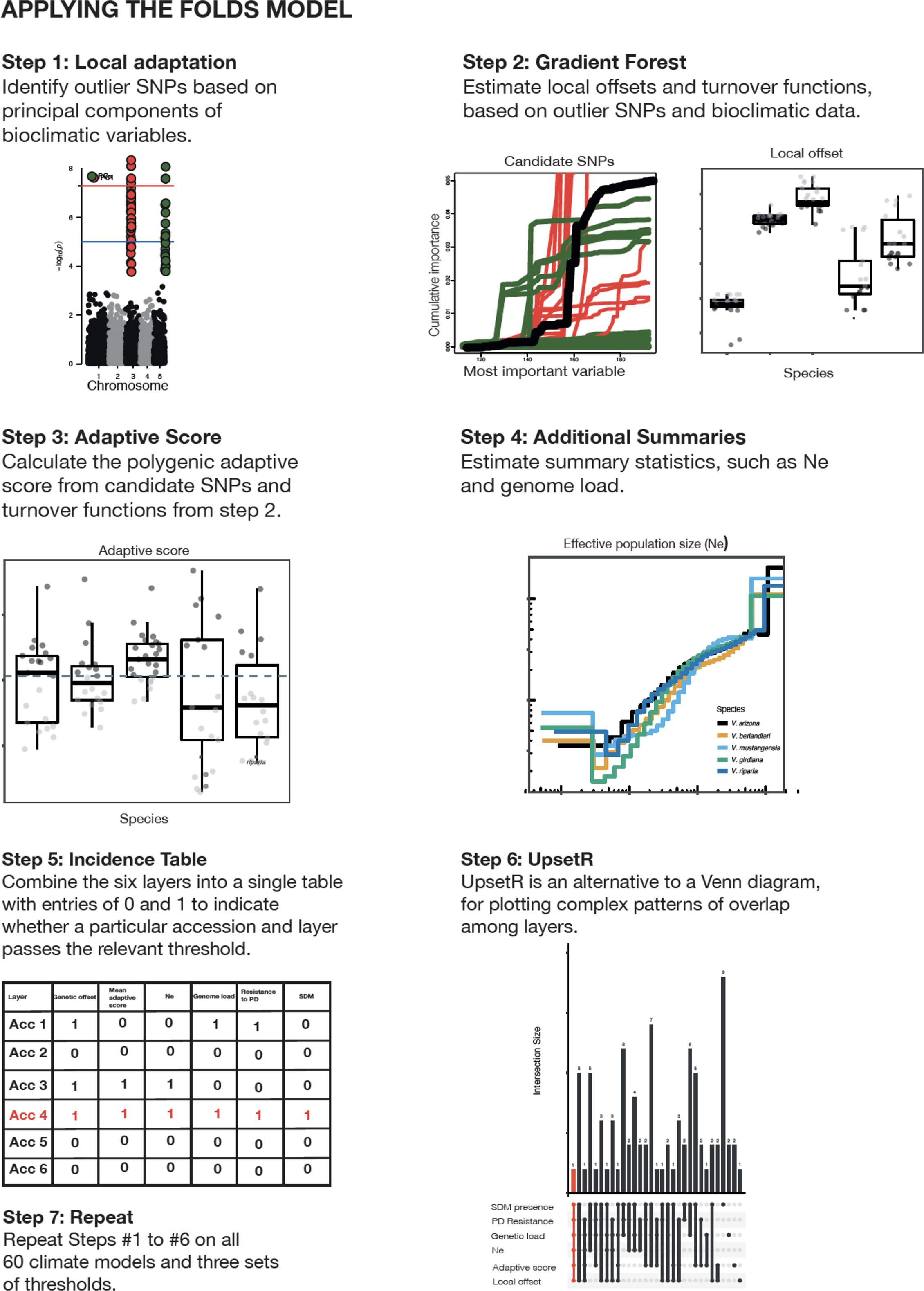
A graphical summary of the steps used in the FOLDS model. Details for each step are provided in the Materials and Methods of the main text.

**Figure S2.**
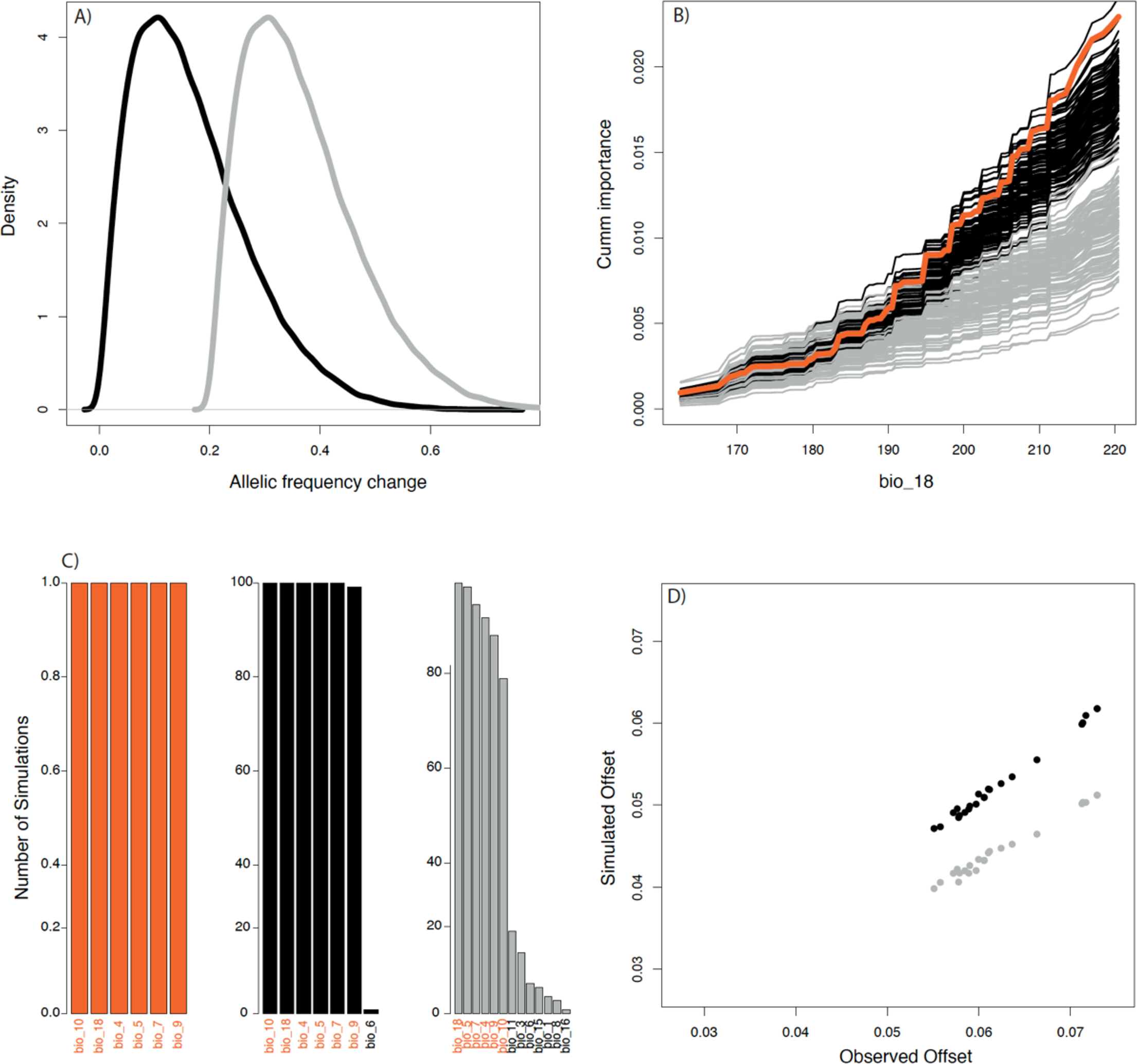
The results of a probabilistic simulation to investigate the effects of using individuals in GF analyses. A) Beta distribution of the magnitude of change of the simulated pseudo-frequencies (Δ) with means 0.1 (black) and 0.3 (grey) for the beta distribution. B) Turnover functions for Bio-18. Black and grey curves show 100 simulations with Δ using a beta distribution with mean 0.1 and 0.3, respectively. The orange curve shows the turnover function for the individual based dataset. C) Histograms showing the number of times the bioclimatic variables were found among the top 6 variables with strong importance (R^2^>0) in the building of the GF models. The orange histogram shows the results obtained with the individual based dataset (1 model). Black and grey histograms show the 100 simulated datasets with different means. The y-axis denoting the proportion of resampled datasets that include the bioclimatic parameter as an important variable. The same bioclimatic variables were routinely identified as important in all three datasets. D) Scatterplot showing genetic offset for one 2070 FCC model of the individual dataset in the x-axis and the mean of the simulated datasets for the different Δ distributions. There is a strong correlation.

**Figure S3.**
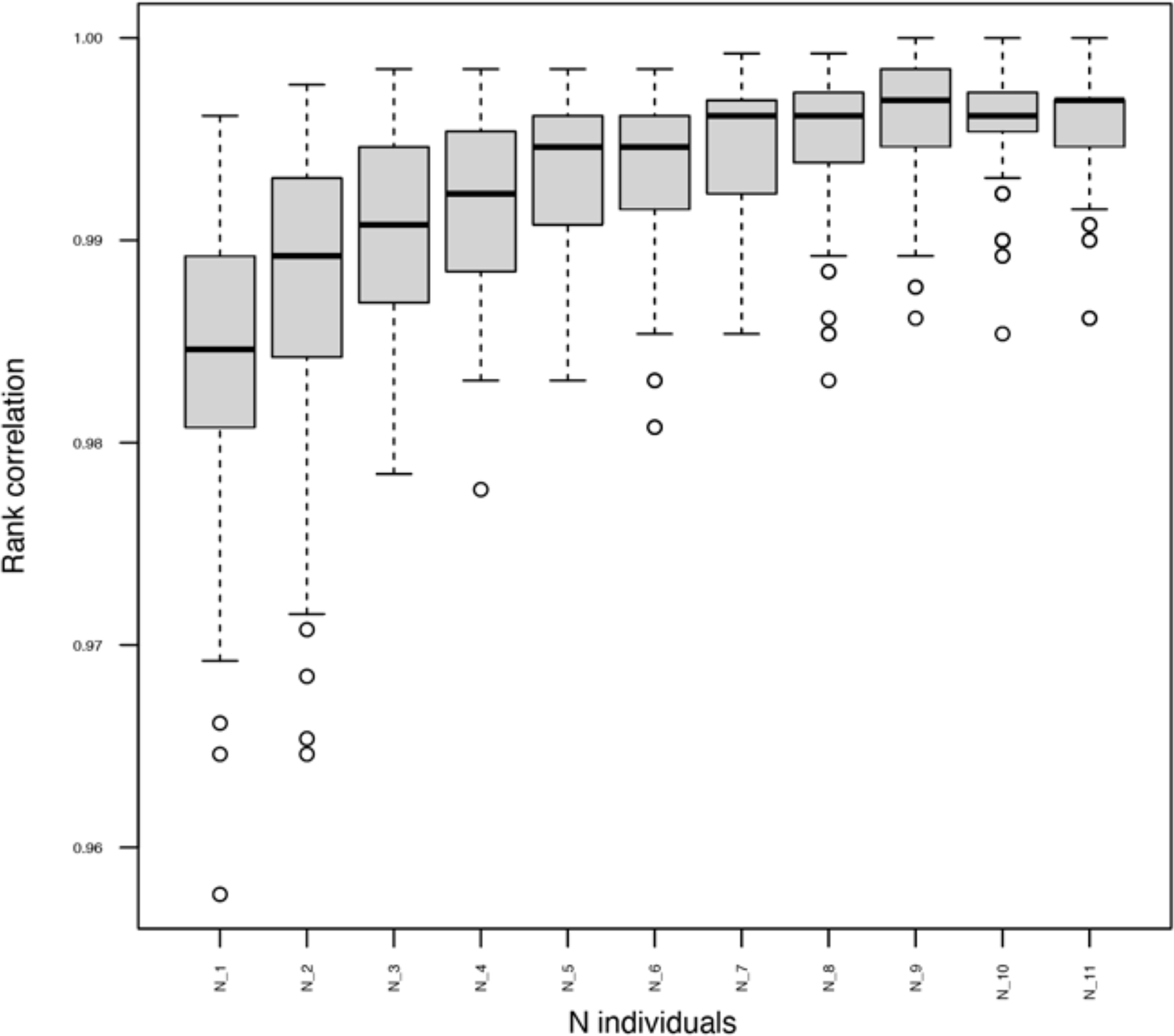
A test to evaluate the effect of subsampling different number of individuals for estimating the genetic offsets across populations, using puplished data of *Zea mays* ssp. *mexicana* (Aguirre-Liguori et al. 2021). Each boxplot shows the rank correlation between the genetic offset across populations subsampling 1 to 11 individuals per population and the genetic offsets using all individuals across all populations. Note that the range of y-axis represents correlations from 0.96 to 1.00. The subsampling were performed 1,000 times per N1 to N11. The results show that for all subsamplings, the rank correlations were >0.96, reflecting that sampling fewer individuals per population – including just one individual - does not strongly affect the order of the genetic offsets across populations.

**Figure S4.**
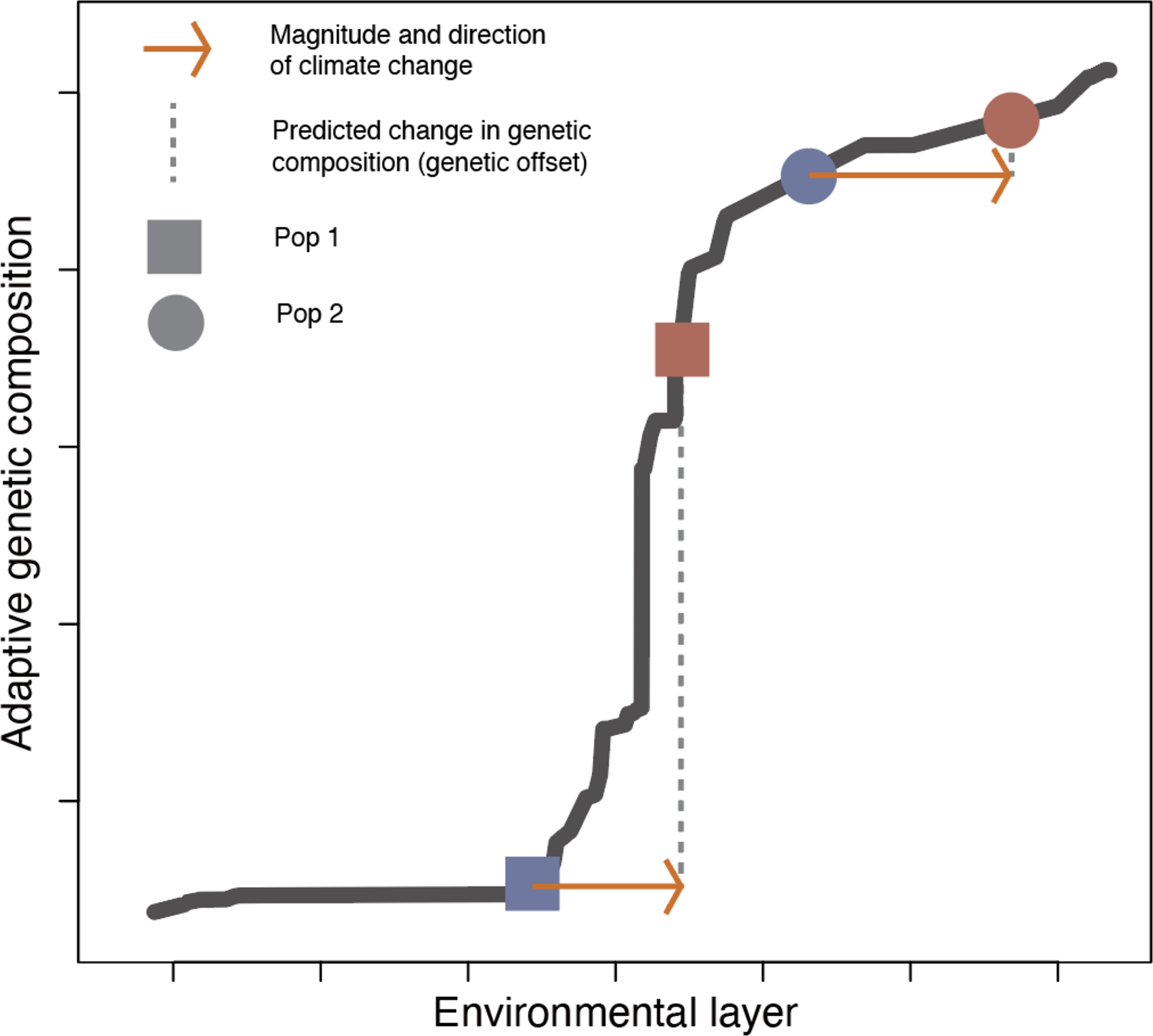
Description of the turnover function and how it can be used to identify the population that is pre-adapted to climate change. The circle and square show two populations along their turnover functions for present (blue) and future (red) climates. The arrows show how climate change will move in the future and the dotted line shows how the genetic composition is expected to change because of climate change (genetic offset). In this scenario, the circled population will have a low offset in the future, and it also grows currently toward the direction where climate will move in the future. Based on this, we consider that from the two populations, the circled one is more pre-adapted to climate change. The importance of this approach is that we infer that pre-adapted populations have the favored allele, which then is included in calculations of *Sa*.

**Figure S5.**
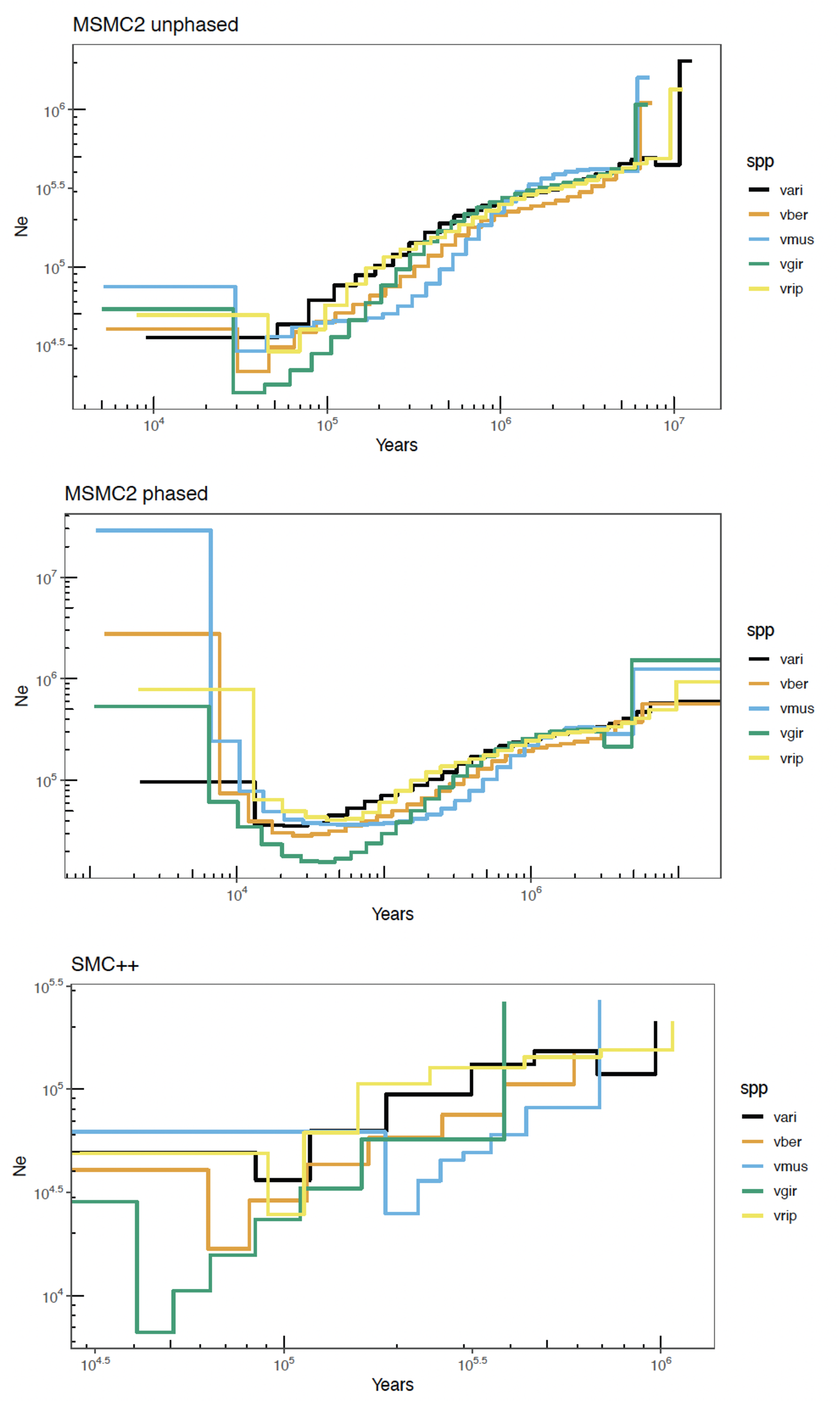
Outcomes from demographic inference. We inferred demographic history based on three different methods: MSMC2 with unphased data (top), MSMC2 with unphased data (middle) and SMC++ (bottom).

**Figure S6:**
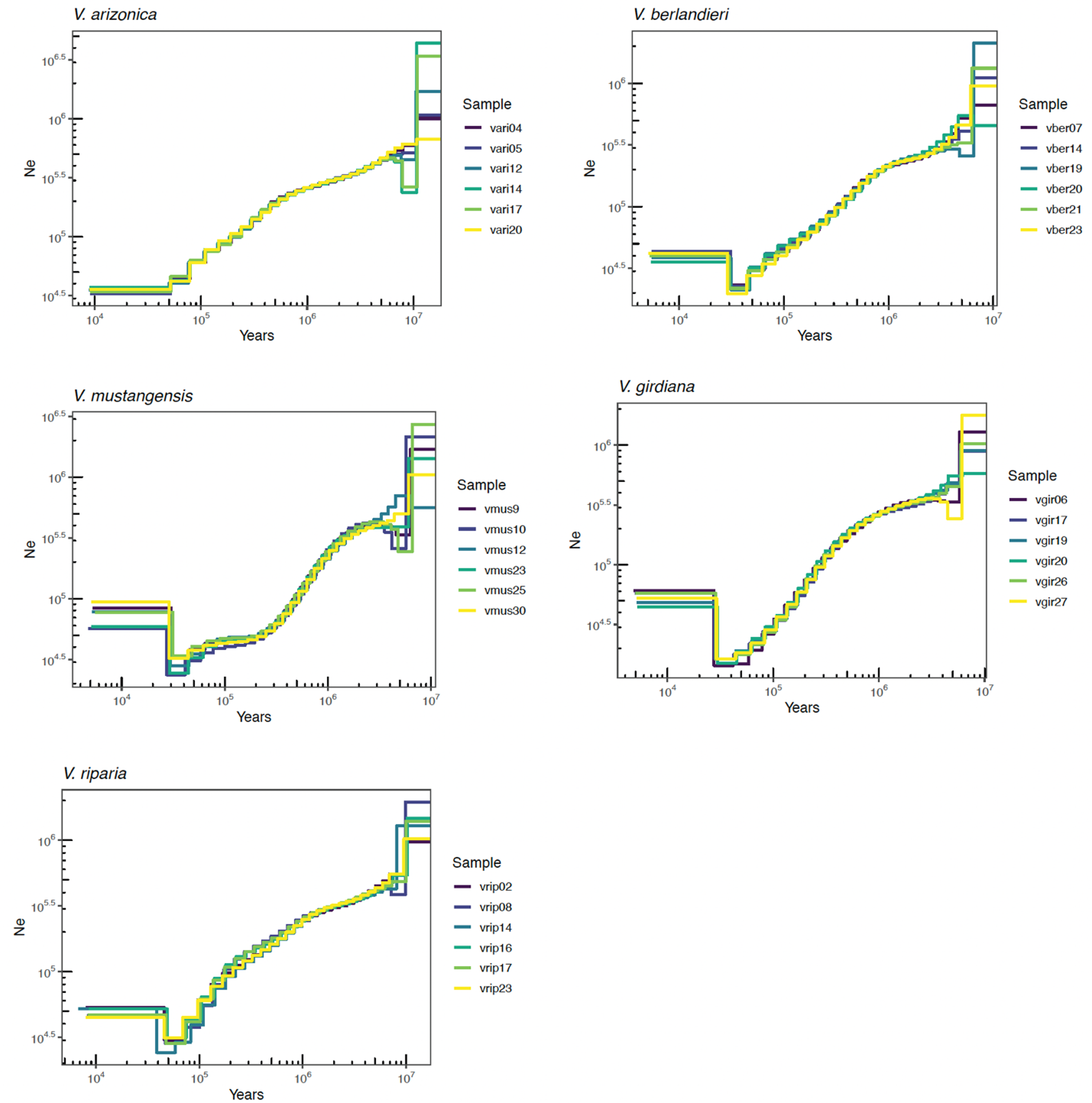
Demographic inference based on individuals in each of the five CWR species. Each graph represents a single species, as labelled, with separate individuals represented by different colors. All inferences were based on MSMC2 with unphased data.

**Figure S7.**
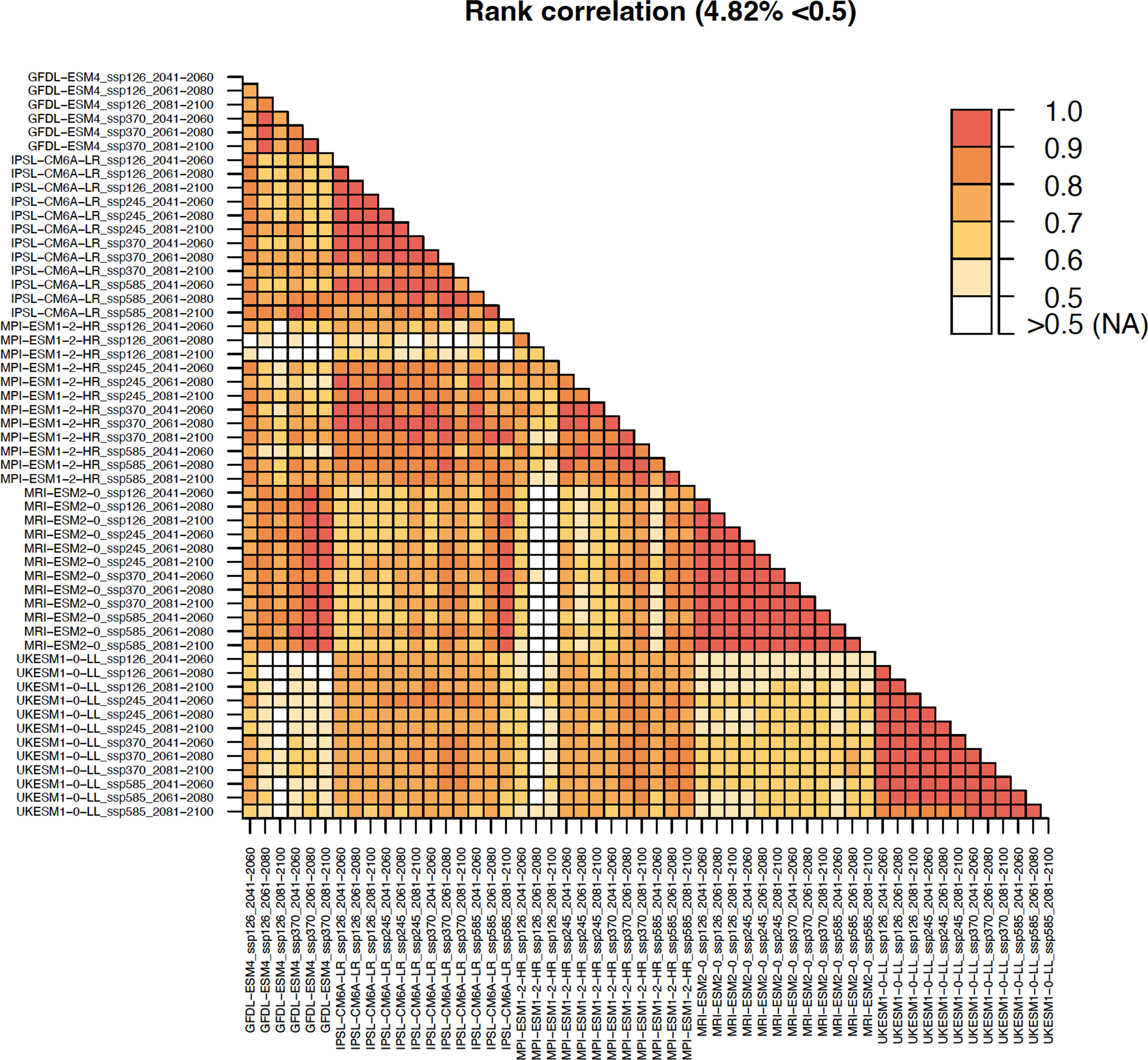
Rank correlations between the genetic offsets of populations using different FCC models. Higher correlations (warmer colors) indicate that the populations have similar offsets between climatic models. White colors indicate that the correlation is below 0.5. We found that only <5% of models were not correlated above 0.5.

**Figure S8.**
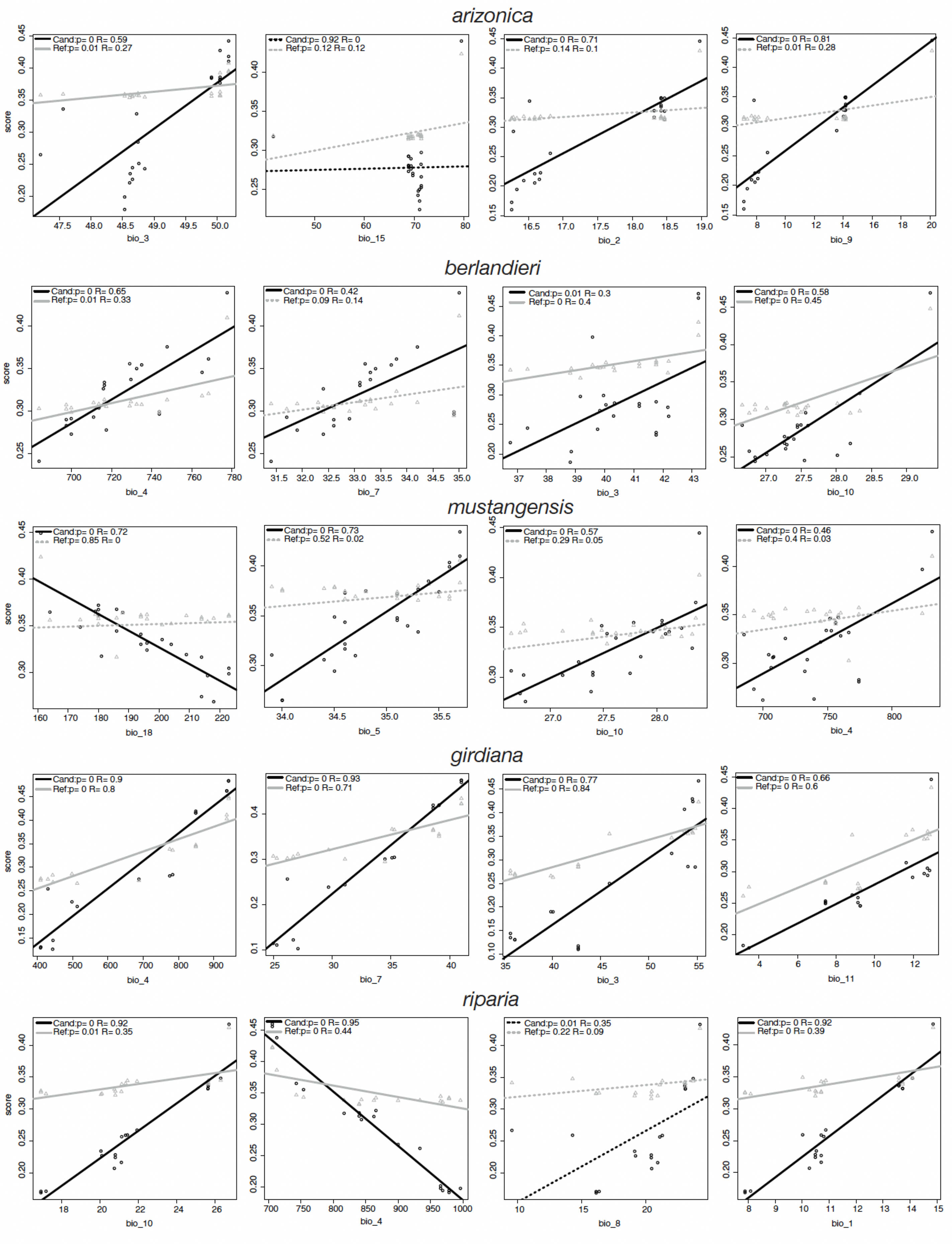
Correlations between adaptive scores and environmental variables. Scatter plots showing the correlation between the top 4 Bioclimatic variables that have a strong contribution to the construction of the GF models (x-axis) and the adaptive scores (y-axis) using the candidate dataset (black, Sa) and the reference dataset (grey, Sref). Continuous lines indicate that the correlation is significant (p<0.05).

**Figure S9.**
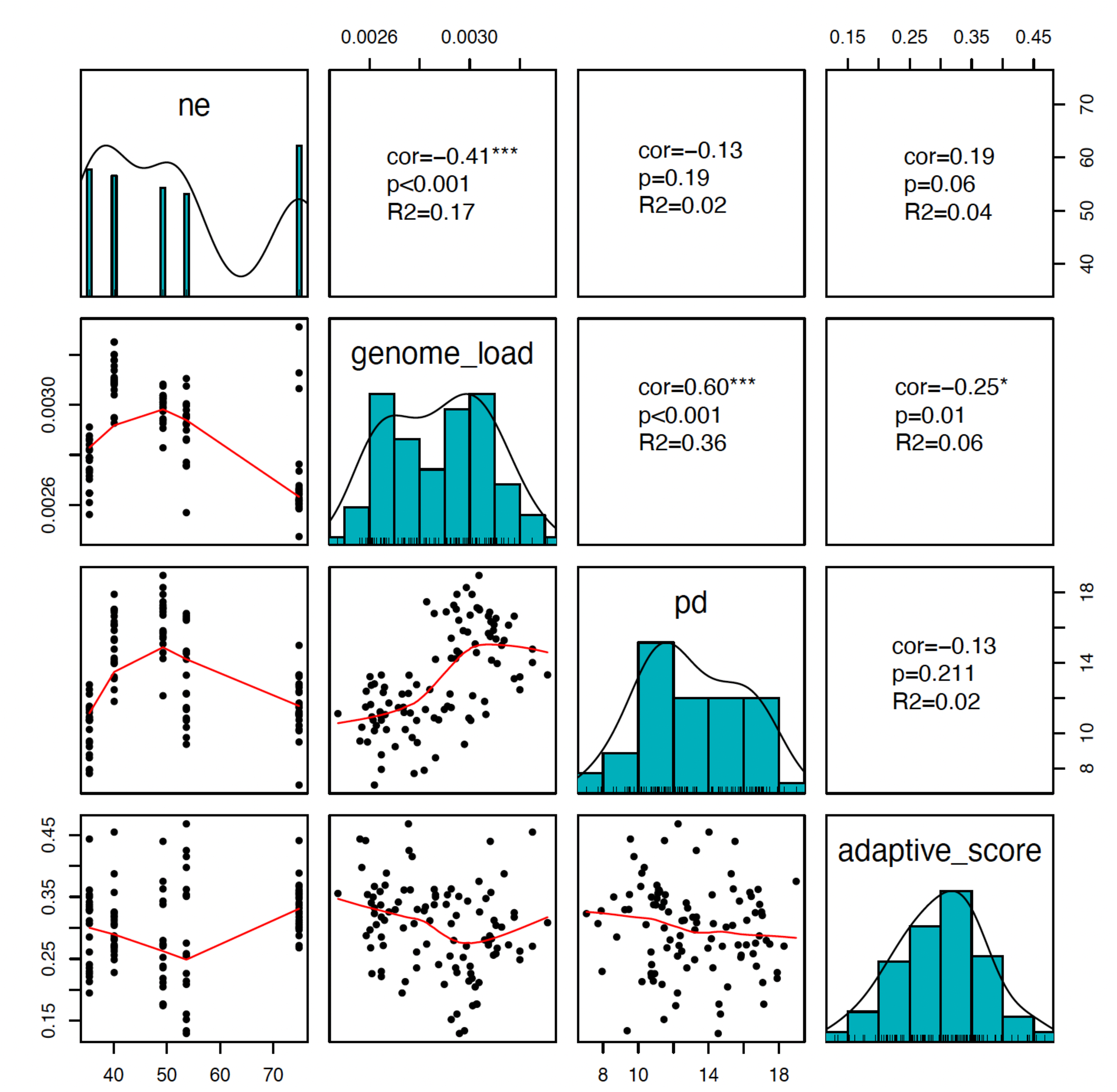
Correlations between variables. Scatterplot matrix showing the correlations between variables across all 105 samples in the study. The diagonal shows the distribution across the samples for each variable. The upper triangle shows the correlations (cor), significance (p) and fit (R^2^) between variables. The lower triangle shows scatterplots for the two sets of variables in question.

### Supplemental tables

**Table S1. Table containing the geographic information, summary statistics and upsetR categories for all the analyses.**

**Table S2.** **Number of SNPs without missing data per species**

| Species | Nº of SNPs |
| --- | --- |
| *Vitis arizonica* | 5,456,474 |
| *Vitis berlandieri* | 3,859,330 |
| *Vitis girdiana* | 3,598,951 |
| *Vitis mustangensis* | 4,971,805 |
| *Vitis riparia* | 5,630,334 |

**Table S3.** **Species distribution models projected to 60 different climatic models. The number indicates the number of 2.5 minute pixels estimated from the raster model.**

| sp | ari | ari | ari | ari | ber | ber | ber | ber | gir | gir | gir | gir | mus | mus | mus | mus | rip | rip | rip | rip |
| --- | --- | --- | --- | --- | --- | --- | --- | --- | --- | --- | --- | --- | --- | --- | --- | --- | --- | --- | --- | --- |
| Year | 2050 | 2070 | 2090 | pres | 2050 | 2070 | 2090 | pres | 2050 | 2070 | 2090 | pres | 2050 | 2070 | 2090 | pres | 2050 | 2070 | 2090 | pres |
| GFDL.ESM4 ssp126 | 54485 | 58518 | 67411 | 51805 | 56468 | 54963 | 57453 | 30029 | 19323 | 20126 | 21529 | 15095 | 70937 | 65132 | 64209 | 19755 | 180616 | 169985 | 166925 | 113839 |
| GFDL.ESM4 ssp370 | 56969 | 61218 | 69913 | 51805 | 79395 | 97830 | 129101 | 30029 | 19436 | 16474 | 18944 | 15095 | 100011 | 146263 | 202641 | 19755 | 196864 | 221273 | 232327 | 113839 |
| IPSL.CM6A.LR ssp126 | 56548 | 57252 | 58406 | 51805 | 80437 | 84458 | 83543 | 30029 | 18866 | 18772 | 18020 | 15095 | 105059 | 106821 | 102604 | 19755 | 199021 | 205215 | 192474 | 113839 |
| IPSL.CM6A.LR ssp245 | 59221 | 65550 | 69563 | 51805 | 92228 | 115360 | 135250 | 30029 | 18419 | 19217 | 18514 | 15095 | 125830 | 168168 | 207271 | 19755 | 210284 | 207974 | 209618 | 113839 |
| IPSL.CM6A.LR ssp370 | 60823 | 67893 | 83174 | 51805 | 98932 | 152333 | 213863 | 30029 | 19223 | 19396 | 18669 | 15095 | 133663 | 229631 | 316748 | 19755 | 208067 | 215631 | 216282 | 113839 |
| IPSL.CM6A.LR ssp585 | 63613 | 75980 | 131522 | 51805 | 107811 | 181300 | 286122 | 30029 | 19891 | 19488 | 17528 | 15095 | 159352 | 282401 | 405209 | 19755 | 214600 | 211546 | 223038 | 113839 |
| MPI.ESM1.2.HR ssp126 | 48393 | 46764 | 46390 | 51805 | 54360 | 53540 | 51109 | 30029 | 18970 | 20190 | 17727 | 15095 | 55562 | 53150 | 59322 | 19755 | 167537 | 161073 | 169915 | 113839 |
| MPI.ESM1.2.HR ssp245 | 47003 | 48431 | 47108 | 51805 | 63428 | 66987 | 77823 | 30029 | 19792 | 18694 | 18406 | 15095 | 74735 | 77700 | 95324 | 19755 | 170836 | 186244 | 200740 | 113839 |
| MPI.ESM1.2.HR ssp370 | 46872 | 50240 | 56193 | 51805 | 66609 | 97229 | 129313 | 30029 | 17764 | 19183 | 18767 | 15095 | 82908 | 142492 | 217062 | 19755 | 193220 | 209029 | 213107 | 113839 |
| MPI.ESM1.2.HR ssp585 | 49419 | 51224 | 62836 | 51805 | 73381 | 105149 | 168470 | 30029 | 18384 | 18209 | 17629 | 15095 | 91240 | 186752 | 276891 | 19755 | 190428 | 216060 | 216682 | 113839 |
| MRI.ESM2.0 ssp126 | 63244 | 62216 | 56194 | 51805 | 72505 | 71338 | 65541 | 30029 | 15504 | 15509 | 15944 | 15095 | 90187 | 85932 | 75711 | 19755 | 199367 | 200138 | 198048 | 113839 |
| MRI.ESM2.0 ssp245 | 61273 | 67748 | 65164 | 51805 | 80997 | 88103 | 85485 | 30029 | 15084 | 15566 | 15097 | 15095 | 105616 | 119560 | 119361 | 19755 | 200003 | 190123 | 204767 | 113839 |
| MRI.ESM2.0 ssp370 | 62961 | 73352 | 85255 | 51805 | 86352 | 102518 | 123293 | 30029 | 15445 | 15069 | 13988 | 15095 | 117808 | 145807 | 180052 | 19755 | 200162 | 209022 | 209300 | 113839 |
| MRI.ESM2.0 ssp585 | 69086 | 80980 | 98989 | 51805 | 95279 | 117911 | 157513 | 30029 | 15147 | 13738 | 15019 | 15095 | 133227 | 170705 | 238317 | 19755 | 205155 | 208343 | 211722 | 113839 |
| UKESM1.0.LL ssp126 | 78063 | 80285 | 81850 | 51805 | 106447 | 117678 | 122283 | 30029 | 17442 | 17674 | 18445 | 15095 | 172072 | 180072 | 184207 | 19755 | 207764 | 203306 | 204821 | 113839 |
| UKESM1.0.LL ssp245 | 81863 | 91580 | 99449 | 51805 | 128707 | 155980 | 180361 | 30029 | 17322 | 19197 | 17761 | 15095 | 199045 | 246452 | 297574 | 19755 | 213758 | 207142 | 199803 | 113839 |
| UKESM1.0.LL ssp370 | 85063 | 99814 | 136035 | 51805 | 147855 | 199399 | 294411 | 30029 | 15475 | 15532 | 15486 | 15095 | 233745 | 335846 | 438705 | 19755 | 213096 | 207323 | 199326 | 113839 |
| UKESM1.0.LL ssp585 | 88831 | 112125 | 172534 | 51805 | 161286 | 238264 | 374755 | 30029 | 17127 | 15319 | 12543 | 15095 | 260358 | 390313 | 478728 | 19755 | 209513 | 200856 | 196519 | 113839 |
| means | 62985 | 69509 | 82665 | 51805 | 91804 | 116685 | 151982 | 30029 | 17700 | 17630 | 17223 | 15095 | 128408 | 174066 | 219996 | 19755 | 198905 | 201682 | 203634 | 113839 |

**Table S4.** **Example showing the table used to create the upsetR analysis. The table shows the summary statitics, and then the future categories for each layer and for each population, based on on the 0.5 threhold. Here we are showing an example using the IPSL.CM6A.LR_ssp585_2061.2080 model The names of the columns indicate the threshold and the layer. In the excel table the genetic offsets and categories are found for all the climatic models and for all the thresholds (names in the columns start with thresh_XX to indicate the threshold).**

|  | spp | lon | Lat | Adaptive score | Genoma load | pd | ne | Mxt IPSL.CM6A.LR ssp585 2061.2080 | Thresh 0.25 adaptive_score | thresh_0.25 genome_load | thresh_pd | thresh_0.25 Ne | thresh_0.25 offset PSL.CM6A.LR ssp585_2061.2080 |
| --- | --- | --- | --- | --- | --- | --- | --- | --- | --- | --- | --- | --- | --- |
| **vrip15** | ari | -111.77 | 34.52 | 0.31 | 0 | 12.49 | 35.45 | 1 | 0 | 0 | 1 | 0 | 1 |
| **vari05** | ari | -108.2 | 32.93 | 0.21 | 0 | 10.21 | 35.45 | 1 | 0 | 0 | 1 | 0 | 1 |
| **vber02** | ber | -99.77 | 30.49 | 0.28 | 0 | 14.16 | 40.11 | 0 | 0 | 0 | 0 | 0 | 0 |
| **vber14** | ber | -98.35 | 30.44 | 0.3 | 0 | 15.36 | 40.11 | 1 | 0 | 0 | 0 | 0 | 0 |
| **vmus15** | mus | -99.14 | 29.35 | 0.35 | 0 | 10.75 | 74.84 | 0 | 1 | 1 | 1 | 1 | 0 |
| **vmus23** | mus | -97.94 | 30.01 | 0.29 | 0 | 12.39 | 74.84 | 1 | 0 | 1 | 1 | 1 | 1 |
| **vgir25** | gir | -116.32 | 36.42 | 0.35 | 0 | 14.2 | 53.67 | 1 | 1 | 0 | 0 | 1 | 0 |
| **vgir10** | gir | -114.43 | 36.39 | 0.47 | 0 | 12.26 | 53.67 | 1 | 1 | 0 | 1 | 1 | 0 |
| **vrip06** | rip | -105.33 | 35.27 | 0.24 | 0 | 14.24 | 49.28 | 0 | 0 | 0 | 0 | 0 | 0 |
| **vrip02** | rip | -104.28 | 37.11 | 0.27 | 0 | 15.68 | 49.28 | 0 | 0 | 0 | 0 | 0 | 0 |

**Table S5. Migration load projected from all accessions defined by the FOLDS test across 54 FCC models to all the vinifera locations in the US (lon and lat). The names of the columns indicate the threshold, the FCC model and the accession used. The column with @cat@ always shows the category 1 and 0, which indicates that the accession could be used or not in the future since they have a migration load below or above the species offset, respectively. This table is in the excel file.**

**Table S6.** **Example showing the migration load projected from a accesion to all the vinifera locations in the US (lon and lat). . The table shows the migration load of the accesion using the IPSL.CM6A.LR_ssp585_2061.2080 model . The names of the columns indicate the threshold, the FCC model and the accession used. The second column always shows the category 1 and 0, which indicates that the accession could be used or not in the future since they have a migration laod below or above the species offset, respectively.**

| lon | lat | tresh_0.25.vmus16.IPSL.CM6A.LR_ssp585_2061.2080 | tresh_0.25.vmus16.cat.IPSL.CM6A.LR_ssp585_2061.2080 | tresh_0.5.vmus16.IPSL.CM6A.LR_ssp585_2061.2080 | tresh_0.5.vmus16.cat.IPSL.CM6A.LR_ssp585_2061.2080 | tresh_0.9.vmus16.IPSL.CM6A.LR_ssp585_2061.2080 | tresh_0.9.vmus16.cat.IPSL.CM6A.LR_ssp585_2061.2080 |
| --- | --- | --- | --- | --- | --- | --- | --- |
| -79.64 | 43.61 | 0.06 | 1 | 0.06 | 1 | 0.06 | 1 |
| -73.73 | 40.78 | 0.06 | 1 | 0.06 | 1 | 0.06 | 1 |
| -77.85 | 40.81 | 0.06 | 1 | 0.06 | 1 | 0.06 | 1 |
| -77.64 | 43.07 | 0.05 | 1 | 0.05 | 1 | 0.05 | 1 |
| -75.9 | 41.26 | 0.06 | 1 | 0.06 | 1 | 0.06 | 1 |
| -119.55 | 34.41 | 0.12 | 0 | 0.12 | 0 | 0.12 | 0 |
| -76.15 | 43.19 | 0.08 | 0 | 0.08 | 0 | 0.08 | 0 |
| -117.17 | 33.63 | 0.1 | 0 | 0.1 | 0 | 0.1 | 0 |
| -121.99 | 37.32 | 0.09 | 0 | 0.09 | 0 | 0.09 | 0 |
| -119 | 37 | 0.1 | 0 | 0.1 | 0 | 0.1 | 0 |

**Table S7. Number of times that an accession is identified by the FOLDS test for 54 FCC models and 3 different thresholds. This table is in the excel file.**

